# Quantifying Support for Competing Biomedical Hypotheses using Large Language Models and Bayesian Analysis

**DOI:** 10.64898/2026.06.05.730173

**Authors:** Bethany M. Moore, Jack Freeman, Robert J. Millikin, Chitrasen Mohanty, Kevin Shine George, Aviral Bal, Cannon Lock, John-Demian Sauer, Megan E. Spurgeon, Darcie L. Moore, Brittany G. Travers, Ron Stewart

## Abstract

**Background:** Comparing competing biomedical hypotheses can help researchers prioritize questions for further investigation. Literature-based evaluation systems such as SKiM-GPT assess support for individual hypotheses, but independently assigned scores may not clearly distinguish their relative support. Direct comparison, accompanied by uncertainty estimates, could help researchers interpret the balance of evidence and track how it changes over time.

**Results:** We introduce KM-GPT-DCH, an extension of SKiM-GPT that integrates co-occurrence-based literature retrieval with large language models to directly compare support for two user-defined biomedical hypotheses. The system evaluates sampled sets of relevant PubMed abstracts using a structured scoring approach and applies a Bayesian model to estimate relative literature support and its uncertainty. In retrospective analyses of five historical hypothesis pairs, KM-GPT-DCH identifies support for subsequently accepted explanations using literature published many years before selected acceptance or rejection milestones. We also apply the system to twenty unresolved hypothesis pairs to characterize literature support and inform future research. Outputs include comparative scores, credible intervals, and explanations linked to retrieved PubMed records.

**Conclusions:** KM-GPT-DCH supports the comparison of biomedical hypotheses and the prioritization of candidates for further investigation, such as, “is HPV or HSV more likely to cause cervical cancer?”. Its temporal analyses enable users to assess and visualize changes in relative literature support. These estimates describe the relative support of literary evidence assessed by the system. A web interface is freely available at https://skim.morgridge.org/.

## Background

Gathering and assessing evidence for and against a hypothesis is central to the scientific method. Researchers integrate findings from published studies and new experiments with domain knowledge to assess competing hypotheses. This process can be understood through a Bayesian perspective, in which prior beliefs are updated according to how probable the observed evidence is under each hypothesis, yielding posterior beliefs [1]. However, the continued, accelerated growth of the scientific literature [2] and computational hypothesis-generation methods makes it difficult for manual evaluation and comprehension in the field even by domain experts [3–15]. Additionally, researchers often need to compare competing hypotheses, which requires careful examination of available evidence, and debate may persist for years or even decades before a conclusion becomes generally accepted or rejected [16].

Research on both large language model (LLM) and human-based evaluation suggests that pairwise comparisons, rather than evaluation in isolation, can improve discrimination between candidates, and in the case of LLMs, align better with human judgement [17–23]. While many computational methods for hypothesis evaluation exist [24–31], including LLM chat interfaces, they are not set up to evaluate hypotheses in a comparative manner, and lack comprehensive scoring and confidence estimation. Additionally, LLM chat interfaces suffer from other limitations like fabrication of information, use of misinformation, and lack of transparency [32–34]. By considering evidence relevant to both hypotheses within a shared evaluation, a comparative approach can assess their relative support rather than relying on separately assigned scores. Such comparisons need not imply that the hypotheses are mutually exclusive or that they exhaust the possible explanations.

Quantifying uncertainty in these comparisons presents an additional challenge. Existing approaches to estimating LLM confidence include verbalized confidence, token probabilities, consistency across responses, trained probes, and Bayesian methods [35]. These approaches address different sources of uncertainty, and some require access to model internals that may be unavailable (see **Supplementary Discussion**). Uncertainty estimations for literature-based hypothesis comparison must account for available literature evidence and variability in the model’s evaluations. Additionally, historical analysis also requires control over the publication dates of retrieved documents. Towards this, our previous retrieval-augmented generation (RAG)-based system SKIM-GPT explored evaluation using time-sliced evidence [36], however tracking relative support for competing hypotheses requires extending this approach to comparative scores and their uncertainty. Restricting retrieval by publication date controls the supplied evidence but does not by itself establish the absence of influence from newer information encoded in the LLM’s parameters (“parametric memory leakage”).

Here, we introduce KinderMiner and Generative Pretrained Transformers with Direct Comparison of Hypotheses (KM-GPT-DCH), a system that extends SKiM-GPT to directly compare literature support for two user-defined hypotheses. The system combines co-occurrence-based retrieval of PubMed abstracts and relevance filtering with standardized comparative prompts and structured outputs, which greatly reduces parametric memory leakage [36]. Repeated LLM evaluations of sampled abstract sets produce evidence-weighted comparative scores. A Bayesian model incorporates these scores, the number of unique supporting abstracts, and an estimate of LLM scoring variability to produce a posterior estimate of relative literature support and a credible interval. These outputs characterize relative literature support under the model’s evaluations and applying the workflow across specified publication windows enables users to examine changes in relative support over time.

We evaluate KM-GPT-DCH using five pairs of hypotheses associated with resolved historical controversies (**Table 1**) and compare the results with co-occurrence-based analysis and independent hypothesis evaluation. These retrospective analyses assess whether the system favors subsequently accepted explanations using literature available at times before scientific consensus occurred, and in all five cases, KM-GPT-DCH determines the correct hypothesis 3 to 12 years before the consensus date. KM-GPT-GCH ensures that the reasoning is based on the retrieved text and not parametric knowledge. We also evaluate the tested conversational LLM interfaces and an agentic system on citation validity and adherence to historical publication cutoffs. Finally, we apply KM-GPT-DCH to twenty unresolved hypothesis pairs to illustrate its use in assessing current literature support and informing further investigation.

**Table 1.**
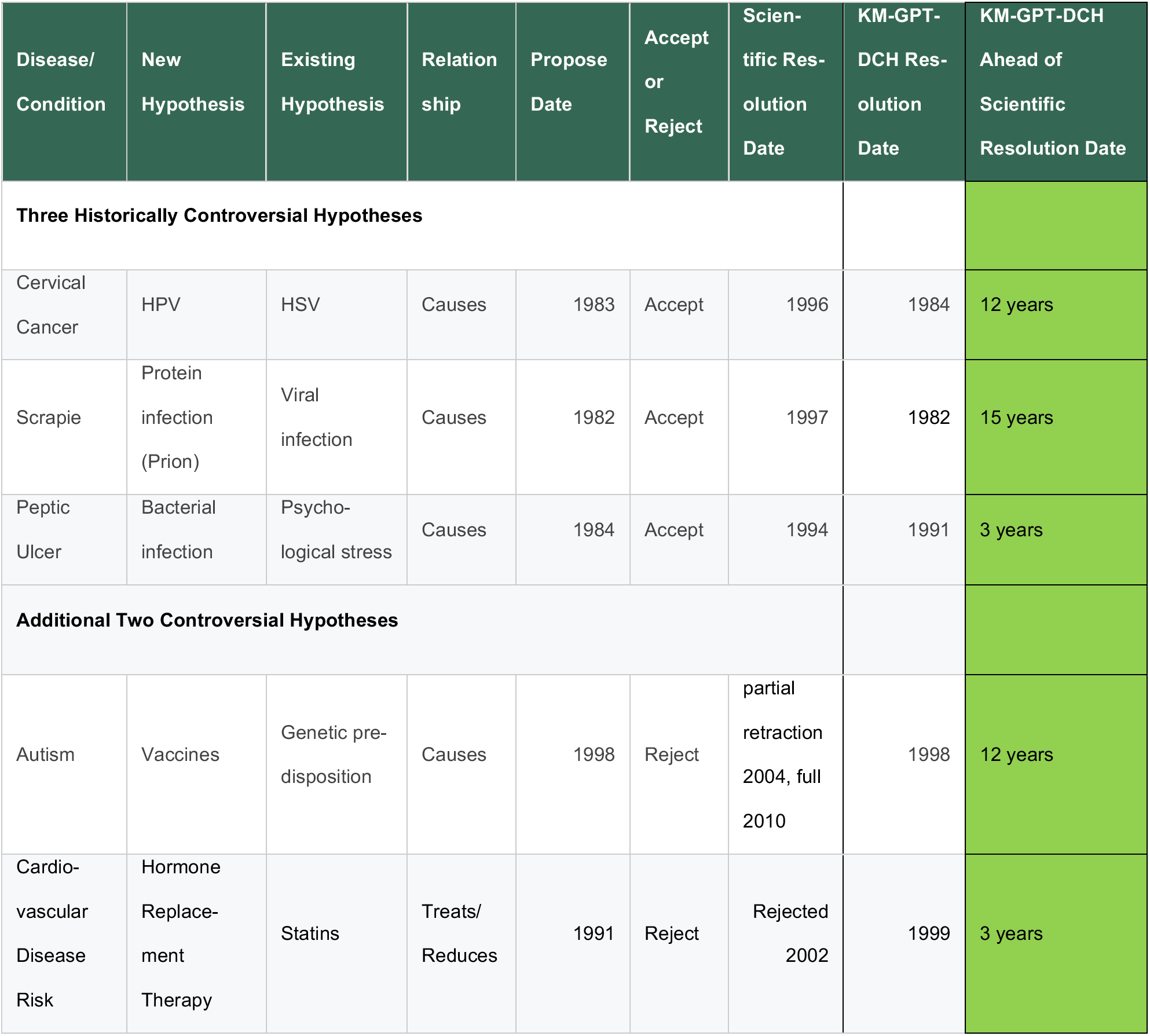
Table of Controversial Hypotheses that are tested using KM-GPT-DCH.

| Disease/ Condition | New Hypothesis | Existing Hypothesis | Relationship | Propose Date | Accept or Reject | Scien-tific Res-olution Date | KM-GPT-DCH Res-olution Date | KM-GPT-DCH Ahead of Scientific Resolution Date |
|---|---|---|---|---|---|---|---|---|
| **Three Historically Controversial Hypotheses** |  |  |  |  |  |  |  |  |
| Cervical Cancer | HPV | HSV | Causes | 1983 | Accept | 1996 | 1984 | 12 years |
| Scrapie | Protein infection (Prion) | Viral infection | Causes | 1982 | Accept | 1997 | 1982 | 15 years |
| Peptic Ulcer | Bacterial infection | Psycho-logical stress | Causes | 1984 | Accept | 1994 | 1991 | 3 years |
| **Additional Two Controversial Hypotheses** |  |  |  |  |  |  |  |  |
| Autism | Vaccines | Genetic pre-disposition | Causes | 1998 | Reject | partial retraction 2004, full 2010 | 1998 | 12 years |
| Cardio-vascular Disease Risk | Hormone Replace-ment Therapy | Statins | Treats/ Reduces | 1991 | Reject | Rejected 2002 | 1999 | 3 years |

KM-GPT-DCH is available through a web interface at https://skim.morgridge.org and an open-source Python package at https://github.com/stewart-lab/skimgpt. The system provides explanations linked to retrieved PubMed records, enabling users to inspect the evidence cited in each evaluation. Its comparative scores, uncertainty estimates, and temporal visualizations support structured assessment of competing hypotheses and prioritization of questions for further research.

## Implementation

### The KM-GPT-DCH algorithm

KM-GPT-DCH leverages both a co-occurrence algorithm (KinderMiner “KM” [13, 37]) and generative pretrained transformers (“GPT”) to directly compare two competing hypotheses (“DCH”). Here, we extend SKiM-GPT, our previous algorithm that evaluates single hypotheses [36], to directly compare one hypothesis (H1) to a second (H2). The KM-GPT-DCH algorithm has several steps illustrated in **Fig. 1**, beginning with KinderMiner to obtain PubMed abstracts from co-occurring terms, followed by a relevance filtering step [36], iterative frontier LLM evaluations (GPT-5.6 Terra from OpenAI) of sampled relevant abstracts, and a Bayesian probability model to estimate uncertainty of the LLM evaluations. KM-GPT-DCH can be run across non-overlapping time retrieval windows to capture how the two hypotheses compare over time. See **Supplementary Methods** for prompt and example outputs.

**Figure 1:**
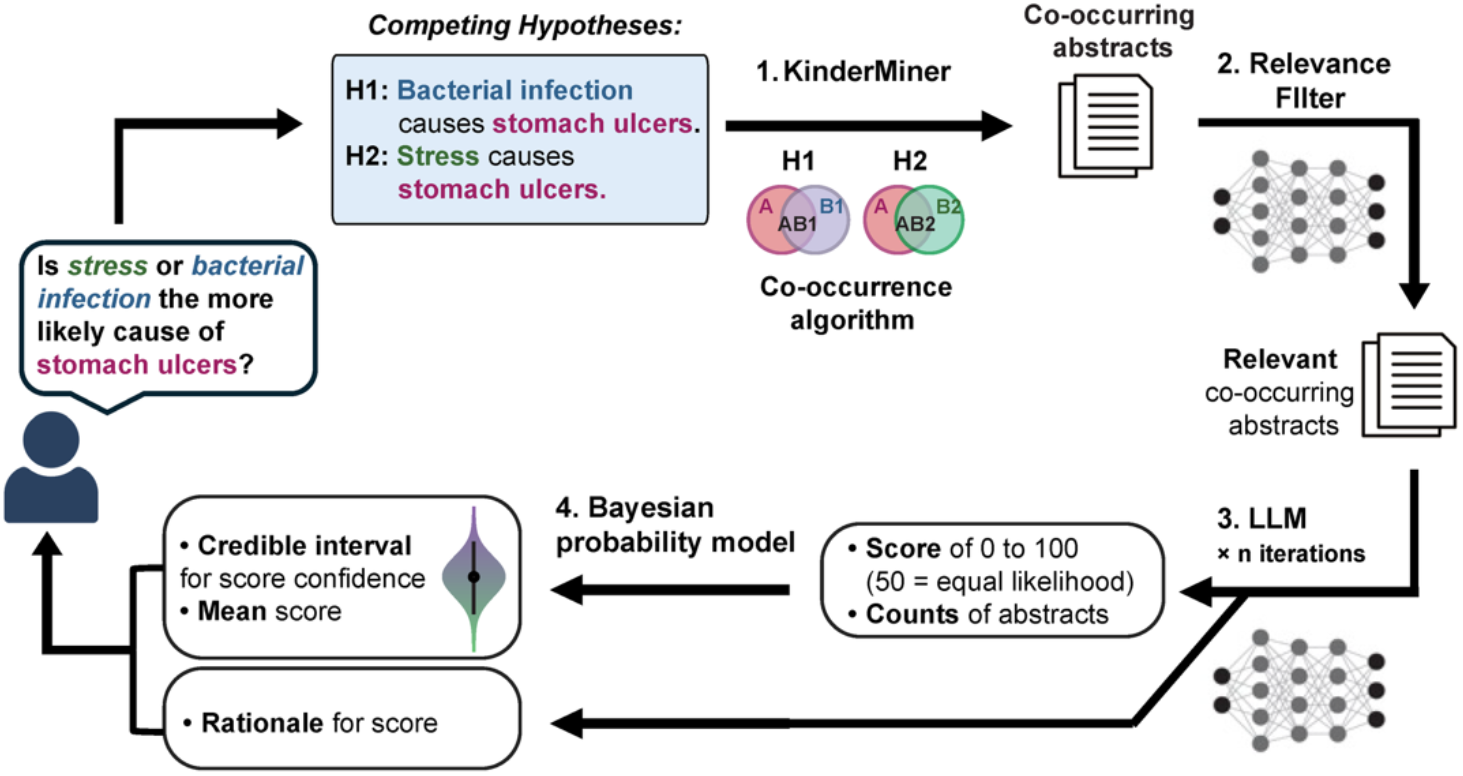
Overview of the KM-GPT-DCH algorithm. 1) **KinderMiner**: the user inputs an A term (e.g., “stomach ulcers”) and two B terms (e.g., “stress” and “bacterial infection”) to construct the two competing hypotheses. Optionally, the user can add a time interval to define the retrieval window for co-occurrence. The co-occurrence algorithm (KinderMiner “KM” [37, 38]) retrieves PubMed abstracts where the A and B terms within a hypothesis co-occur. 2) **Relevance filter.** The abstracts identified from step 1 above are filtered for relevance to each hypothesis with a fine-tuned small language model [36]. 3) **Frontier LLM**. The LLM adjudicates between the two hypotheses using the two relevant literature pools as its sole evidence, a standardized prompt, and output structure. This framework forces a direct comparison of the competing claims. For example, weighing whether stress or bacterial infection is the more probable cause of stomach ulcers requires the model to provide a score and rationale based on the relative strength of the cited retrieved data. The score provided by the LLM ranges from 0 (very strong support for H2) to 100 (very strong support for H1). Here we use the OpenAI GPT-5.6 Terra model. **4) Bayesian probability model**. Finally, a Bayesian credible interval (95% credibility) of the score is calculated (see **LLM score and statistics**). KM-GPT-DCH: KinderMiner and Generative Pretrained Transformers with Direct Comparison of Hypotheses LLM: Large Language Model H1: Hypothesis 1 H2: Hypothesis 2

### KM co-occurrence statistics

We compare co-occurrence results (from KinderMiner, or KM) using the z test of proportions: 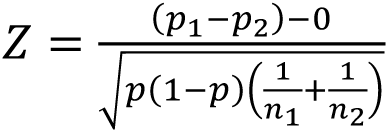 where p1 is proportion 1 or the ratio of AB counts over B counts for B term 1, p2 is proportion 2, or the ratio of AB counts over B counts for B term 2, and p is the overall sample pooled proportion from both terms. n1 and n2 are the sample sizes for the respective groups. The *z*-score was then compared to the *z*-statistic associated with the given alpha level to obtain a *p*-value.

If co-occurring abstracts were only available for one of the two hypotheses, then the *p*-value was set to 1*e*^-300^ (capped) for the hypothesis with abstracts, and 1 for the hypothesis with no co-occurring abstracts. The *z*-statistic was set to the highest value if the hypothesis with the abstracts was the first hypothesis, or the lowest value if the hypothesis with abstracts was the second hypothesis.

### Sample Retrieval and LLM Score

To construct the evidence base for each evaluation, the algorithm retrieves deduplicated PubMed abstracts from a given window of years and applies a stratified random sampling method to construct sets of 50 abstracts. This sampling enforces a minimum sampling floor of 6% from each hypothesis’s literature pool to ensure that both hypothesis pools are represented even when one is much larger than the other. If the total pool of relevant abstracts exceeds 50, the algorithm utilizes sampling with replacement across R = 10 iterations to capture different document sets. If the total pool of relevant abstracts is greater than 500, we set R = 20. For retrieval of abstracts in this paper, a window of 5 years is set, so that for a given year y, abstracts from year y as well as the previous 4 years are retrieved (ie. year 2000 retrieves abstracts from 1996-2000). The user generated hypotheses and abstracts are then input into the LLM and scored where a score of 100 indicates full literature support for H1, and 0 indicates full literature support for H2, and a score of 50 indicates each hypothesis has equal support or no support (see **Supplementary Methods** for this prompt). Each iteration is scored separately by the LLM; the total unique abstracts (*n_lit_*) is calculated across all iterations (after abstracts have passed the relevance filter).

### Statistical Model

The Bayesian statistical model estimates what fraction (or proportion) of the literature corpus supports H1 relative to H2 in time period *t*, given the abstracts retrieved from PubMed. Because H1 and H2 are competing hypotheses, and we wish to estimate bias (literature support, Θ) given the number of trials (abstracts, *n*), we model literature support as a Bernoulli process. The output of this model is a probability distribution for Θ given the retrieved abstracts.

The abstracts are provided to a large language model (LLM) along with instructions to generate a score between 0 and 100; a score of 100 indicates full literature support for H1, and a score of 0 indicates full support for H2 (see **Supplementary Methods** for this prompt). The score is divided by 100 to place it on a 0 to 1 scale, and this value (*x*) is multiplied by the number of abstracts, *n*, to arrive at the number of Bernoulli trials supporting H1, *k*; the number of Bernoulli trials supporting H2 is then *n* − *k* (for example, if the LLM was provided with 10 abstracts and generates a score of 70, then H1 is supported by 0.7 ⋅ 10 = 7 trials and H2 is supported by 10 − 7 = 3 trials). This sampling process can be run *R* times to draw different abstract sets from the literature corpus; in this case, the LLM scores are averaged, and *n* is the total number of unique abstracts passed to the LLM (see **Sample Retrieval and LLM Score** for details). Abstracts that support neither H1 nor H2 are subtracted from *n* before calculating *k*. Rather than simply counting abstracts that support H1, this method is employed so as to allow the LLM to weigh the information contained in the abstracts, in effect producing a weighted vote count according to the LLM’s judgment.

The probability distribution for bias in a Bernoulli process given the observed values of *n* and *k* is given by the Beta distribution (for now, using a uniform Beta(1,1) prior):

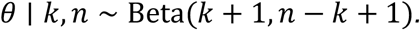

We modify this simple model in two ways. First, we account for the stochasticity in LLM scoring; *i.e.*, the LLM can produce different scores given the same set of abstracts. Second, we modify the Bayesian prior to regularize the model for when there are few or no abstracts.

We estimate the variance due to LLM scoring, *σ_llm_*^2^, by providing the same set of abstracts to the LLM 50 times and calculating the variance of the generated scores; this process is repeated with 20 different sets of hypotheses and their abstracts. The resulting data indicate that this variance depends on *θ* (*σ_llm_*^2^ tends to be lower if *θ* is close to 0 or 1, consistent with the Bhatia-Davis inequality [39]; we approximate *σ_llm_*^2^ ≈ 0.0076 ⋅ *θ*(1 − *θ*) via linear regression (see **Supplementary Methods**). Given *R* abstract sampling iterations, with each iteration’s abstracts scored by the LLM independently, the variance of the mean score due to LLM scoring is *σ_llm_*^2^/*R*.

Assuming these two sources of variance are independent, the total variance in the mean score is the sum of the variance from *n* Bernoulli trials and the variance from LLM scoring:

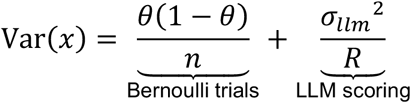

The variance from LLM scoring must be expressed in units of Bernoulli trials before it can be used in a Beta model. We refer to *n_eff_* as the effective number of Bernoulli trials that would produce this total variance, and substitute the approximation for *σ_llm_*^2^ obtained earlier via linear regression:

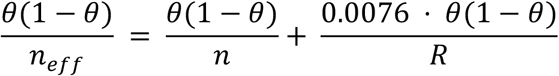

The *θ*(1 − *θ*) terms cancel, leaving:

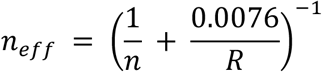

Thus, the number of trials supporting H1 accounting for variance in LLM scoring, *k_eff_*, is *n_eff_* ⋅ *x* where *x* is the LLM score, and the number of trials supporting H2 is *n_eff_* − *k_eff_*. A special case is when *n* = 0, for which we set *n_eff_* = 0.

Lastly, we modify the Bayesian prior, encoded as a Beta distribution, *θ* ∼ Beta(*a*_0_, *a*_0_), which is equivalent to already having observed *a_0_* hypothetical trials supporting H1 and *a_0_* trials supporting H2. By default, we use *a_0_* = 1.2 so the posterior is guaranteed to be unimodal and slightly shrunk towards *θ* = 0.5. Because the Beta prior is conjugate to the Bernoulli likelihood, *a_0_* can simply be added to the number of trials supporting each hypothesis to give the posterior:

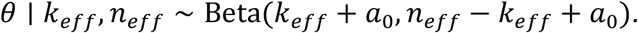

The posterior mode is reported as a point estimate of *8*; this point estimate should be interpreted as how supportive the literature was of H1 relative to H2 in each time period, not the probability that H1 is factually correct. The 95% highest density interval (HDI) is the narrowest credible interval containing 95% of the posterior probability [40] and is reported as an interval estimate of *8*. This interval estimate should be interpreted as the precision with which *8* can be estimated by our pipeline. Generally, a large credible interval width occurs when few relevant abstracts were retrieved for time period *t*.

### Criteria for determining whether a hypothesis pair is “resolved”

We define a historical hypothesis to be resolved when the posterior score is at least 67 (when H1 is approximately twice as supported as H2) and the credible interval does not cross the 50-mark line.

### Finding unresolved hypothesis pairs

To identify additional controversial hypotheses, we used llm-council (GitHub: karpathy/llm-council). Given a query, llm-council orchestrates a panel of LLMs and a Chair; first the query is passed to the panel LLMs individually, and the responses are collected and anonymized responses are passed to other panel LLMs to rank them in accuracy and insight. The Chair collects all the models’ responses and compiles them. We provided our existing list of curated hypotheses to llm-council and asked to generate a newer set of unresolved hypothesis pairs (see **Supplementary Table 4**). The collected hypotheses were subsequently reviewed manually.

## Results

### Competing historical hypotheses

We begin by evaluating three sets of historical hypotheses to provide a proof of concept for the KM-GPT-DCH algorithm and to compare with other methods: 1) bacterial infection causes peptic ulcers [41, 42], 2) human papillomavirus (HPV) causes cervical cancer [43–45], and 3) prions cause scrapie [16, 46]. Each set has a clear pre-existing hypothesis that was subsequently supplanted by a new hypothesis that took approximately a decade to be accepted. We then evaluate two additional sets of hypotheses in which one of the hypotheses is subsequently refuted after several years: 1) Autism is caused by genetics not vaccines [47] and 2) Cardiovascular disease (CVD) can be better treated by statins not hormones [48, 49]. Controversial hypotheses with their proposal date and acceptance or refutation date by the scientific community are listed in **Table 1**; for full table including public acceptance and refutation dates see **Supplementary Table 1**. See **Supplementary Methods** for how the hypothesis acceptance and refutation dates were determined. In the text, unless otherwise specified, acceptance and refutation dates refer to the scientific community acceptance and refutation dates. General public acceptance/refutation dates typically lag behind the scientific community’s dates.

### Conversational LLM interfaces are insufficient for direct hypothesis comparison

Because of their widespread use and general usability, we first assessed if chat-based LLM interfaces could reliably determine if one hypothesis is better supported than another. We tested five commonly used conversational LLM interfaces and one multi-agent system (Crow) on citation accuracy and whether the correct historical hypothesis could be determined at the appropriate time. For fair analysis of these historical hypotheses, it is necessary to perform temporal filtering (only using references up to a specific date). With temporal filtering, we found that the five conversational LLMs only provided 15% to 25% error-free legitimate references (85% for Crow). All six systems get the correct answer (for the leading hypothesis at a given time) only 67% to 80% of the time. This highlights the unreliability of these systems as well as the inability to be temporally aware. While newer versions of conversational LLMs may have smaller error rates, there is always a hallucinatory element; our RAG system always provides abstracts from real papers. Furthermore, conversational LLM interfaces do not provide a level of confidence in their answer, nor can they perform hypothesis evaluations at scale. For details on the systems, versions, and methods see **Supplementary Discussion;** see **Supplementary Tables 2** and **3** for summarized results and **Supplementary Data 1** for all LLM response results.

### Peptic ulcer hypotheses comparison using KM co-occurrence or KM-GPT or KM-GPT-DCH

Traditionally, peptic, gastric, or duodenal ulcer (“peptic ulcer”) was thought to be caused by psychological stress or other factors [50, 51]. In 1984, the bacterium *Helicobacter pylori* was implicated in peptic ulcer by Warren and Marshall [41, 52]. Bacterial infection was not generally accepted as a primary cause of peptic ulcer until 1994 when the National Institutes of Health came to a consensus [53], with Warren and Marshall later winning the 2005 Nobel prize. (See **Supplementary Discussion.**)

Given that using a straightforward co-occurrence model would be more computationally efficient, we first tried our KM co-occurrence method alone [37] to investigate these competing hypothetical causes of peptic ulcer. In contrast to LLM conversational interfaces, KM always provides real references and can correctly perform time filtering. Using only KM, however, peptic ulcers were not found to have a stronger association with bacterial infection (compared to psychological stress) until 1999, well after the NIH consensus accepted this idea (**Fig. 2a**). We concluded that while KM is useful in reliably finding associated abstracts to review, the calculated *p*-values from co-occurrence alone are not sufficient to evaluate competing hypotheses.

**Figure 2.**
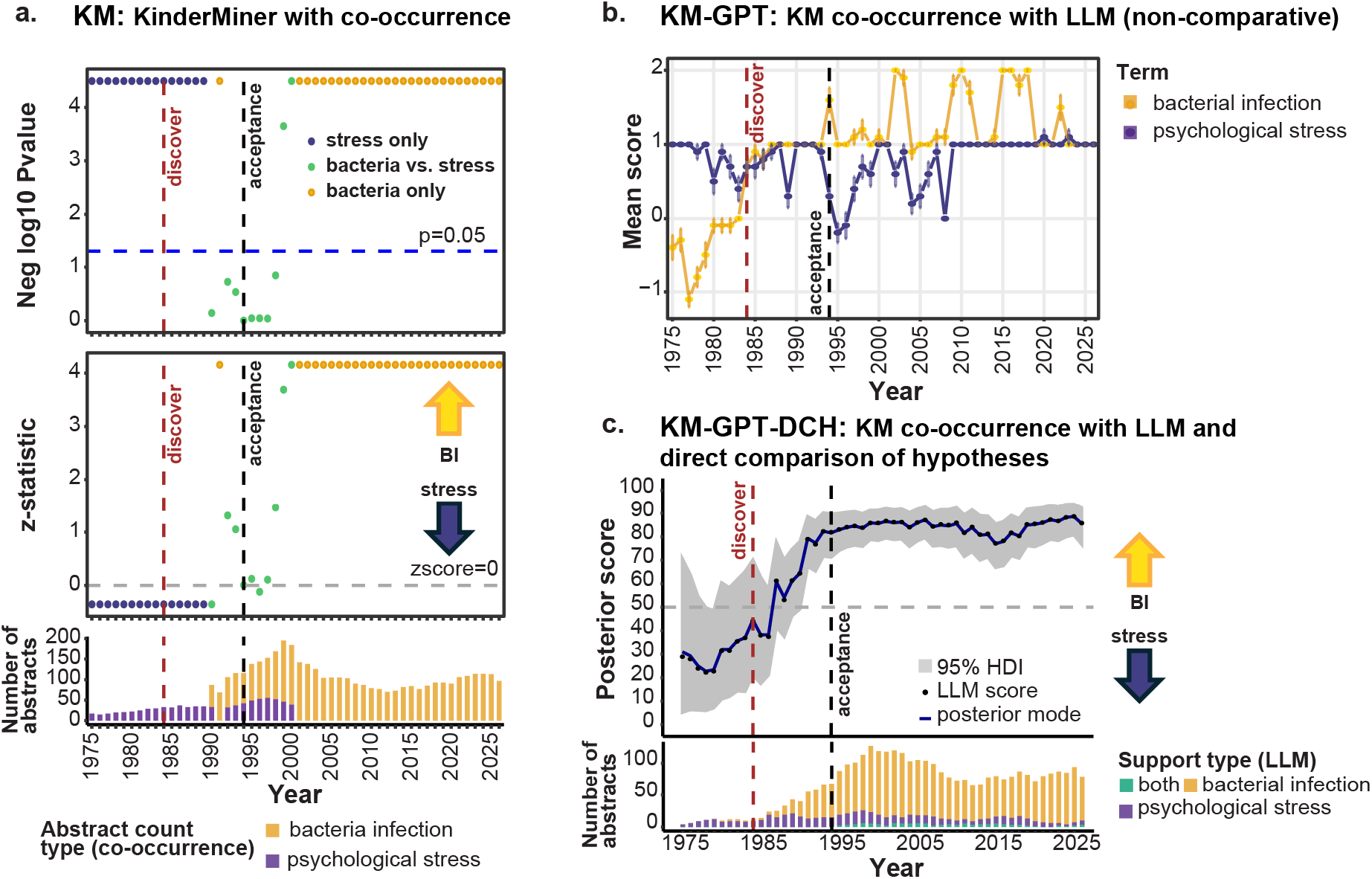
KM, KM-GPT, and KM-GPT-DCH applied to the competing hypotheses for peptic ulcer causation. **a.** KM co-occurrence results for each year between the competing hypotheses for the causation of peptic ulcer: bacterial infection (“BI”) vs. psychological stress (“stress”). The top panel shows the negative log10 of the *p*-value; the middle panel shows the *z*-statistic; the bottom panel shows the number of abstracts obtained for each term. Years range from 1975-2026. In all graphics in Fig. 2, the red vertical line indicates the discovery date for bacterial infection (1984), while the gray line indicates the acceptance date for bacterial infection (1994). **b.** KM-GPT results when evaluating the two peptic ulcer hypotheses independently from 1975-2026, where the score from the OpenAI GPT-5.6 Terra model ranges from-2 to 2, with negative scores refuting the hypothesis and positive scores supporting the hypothesis. **c.** KM-GPT-DCH results. The graph is the mean posterior score from KM-GPT-DCH (iterations=10) where a score above 50 is more supportive of H1 (bacterial infection), a score below 50 is more supportive of H2 (psychological stress), and a score of 50 indicates equivalent or inconclusive support or that no abstracts are available. Only the combined comparative system, KM-GPT-DCH, successfully identifies bacterial infection as the more likely cause of peptic ulcer years before (1991) general acceptance (1994). A Bayesian credible interval (gray shaded region) for each time point is computed and the 5th and 95th percentile of the posterior is shown (see **Methods**). The bar plot at the bottom shows the number of abstracts used for the evaluation each year with the color scale designating the number supporting the first hypothesis, the second hypothesis, or both hypotheses. RAG: retrieval augmented generation LLM: large language model KM: KinderMiner co-occurrence algorithm DCH: direct comparison of hypotheses GPT: generative pre-trained transformer

We next utilized our RAG-based single hypothesis evaluator, KM-GPT [36], to evaluate the hypotheses. When we applied KM-GPT to the two hypothetical causes of peptic ulcer separately (**Fig. 2b**), the hypotheses had similar scores, revealing that issues like score shrinkage make non-comparative analysis untenable [21–23]. In summary, neither the co-occurrence algorithm KM nor KM-GPT was able to provide early resolution of whether bacterial infection or psychological stress is a more likely cause of peptic ulcer.

It is only when we directly compare the hypotheses of psychological stress and bacterial infection that we can effectively determine the more plausible hypothesis for peptic ulcer causation. The KM-GPT-DCH algorithm performs this direct comparison by using KM to retrieve abstracts which are then provided to a frontier LLM to directly compare evidence for the two hypotheses. The algorithm provides a score from 0 to 100, where above 50 is more supportive of H1 (bacterial infection) and below 50 is more supportive of H2 (psychological stress). Repeated LLM evaluations of sampled abstract sets yield comparative scores that are combined with the number of unique supporting abstracts in a Bayesian model to estimate relative literature support and its credible interval (see **Methods**). We applied KM-GPT-DCH over time from 1975-2026 using 5-year windows (i.e. for the year 1980 the window range would be 1976-1980) to the peptic ulcer case (**Fig. 2c**). Before 1985, the score was mostly below 50, indicating stronger support for psychological stress relative to bacterial infection as the cause of peptic ulcer. From 1991 onwards, the score remained at 70 or higher with the credible interval not crossing 50, indicating strong support for the bacterial infection hypothesis relative to the psychological stress hypothesis and demonstrating that KM-GPT-DCH chose the more supported hypothesis before the general acceptance of the bacterial infection hypothesis in 1994. Mild support for psychological stress persisted, indicating that while bacterial infection was the stronger hypothesis, there was modest literature support for psychological stress as a risk or complicating factor in peptic ulcer [51]. We also show that KM-GPT-DCH can evaluate a wider range of papers (up to 120 abstracts for the peptic ulcer comparison, more in other comparisons, **Fig. 2c**) to get a more comprehensive view than other LLM-based systems (which usually evaluate only 11-12, see **Supplementary table 2**). See **Supplementary Discussion** for an example of the response and reasoning for the cause of peptic ulcer in 1987. Note that in this example the LLM adjudicates in favor of the bacterial infection hypothesis even though the number of abstracts is typically greater for the psychological stress hypothesis indicating that the LLM pays attention to the strength of support in each abstract. All KM-GPT-DCH results, reasoning, and scores are in **Supplemental data 2**. The existence of a third cause hypothesized to be a plausible cause for peptic ulcer, NSAIDs, is addressed in the **Discussion**.

### KM-GPT-DCH evaluation of hypotheses about cervical cancer and scrapie

We next directly compare the two hypotheses for the cause of cervical cancer with KM-GPT-DCH run over time from 1975 to 2026. Herpes simplex virus (HSV) was thought to be a potential cause of cervical cancer prior to 1983, as HSV was highly correlated with carcinoma of the cervix [54]. Human papillomavirus (HPV) was proposed to be the cause of cervical cancer in 1983 by Harald zur Hausen and colleagues [43, 55]. However, the HPV hypothesis was not generally accepted until 1996, when the National Institutes of Health came to a consensus on the primary cause of cervical cancer [56]. Dr. zur Hausen won the Nobel Prize in 2008.

The results shown in **Fig. 3a** indicate that while HSV was the leading cause from 1976-1982 with an average score of 37, from 1983 onward, the algorithm favored HPV with average scores between 58 in 1983 to 95 in 2025. In our judgement, the system provides sound reasoning for its scoring based on the abstracts it is given. An average score of 78 in 1984, just one year after the landmark paper was published, shows that KM-GPT-DCH detects what will be the prevailing hypothesis (HPV) 12 years before this is formally accepted as true (in 1996). Additionally, the credible interval, which is based both on the evaluation scores as well as the number of abstracts supporting each hypothesis (see **Methods**), is wider earlier on (1975-1985), indicating more score variability and less abstract support; but this narrows in later years, indicating an increase in both consistent scoring and number of abstracts (**Fig. 3a**).

**Figure 3.**
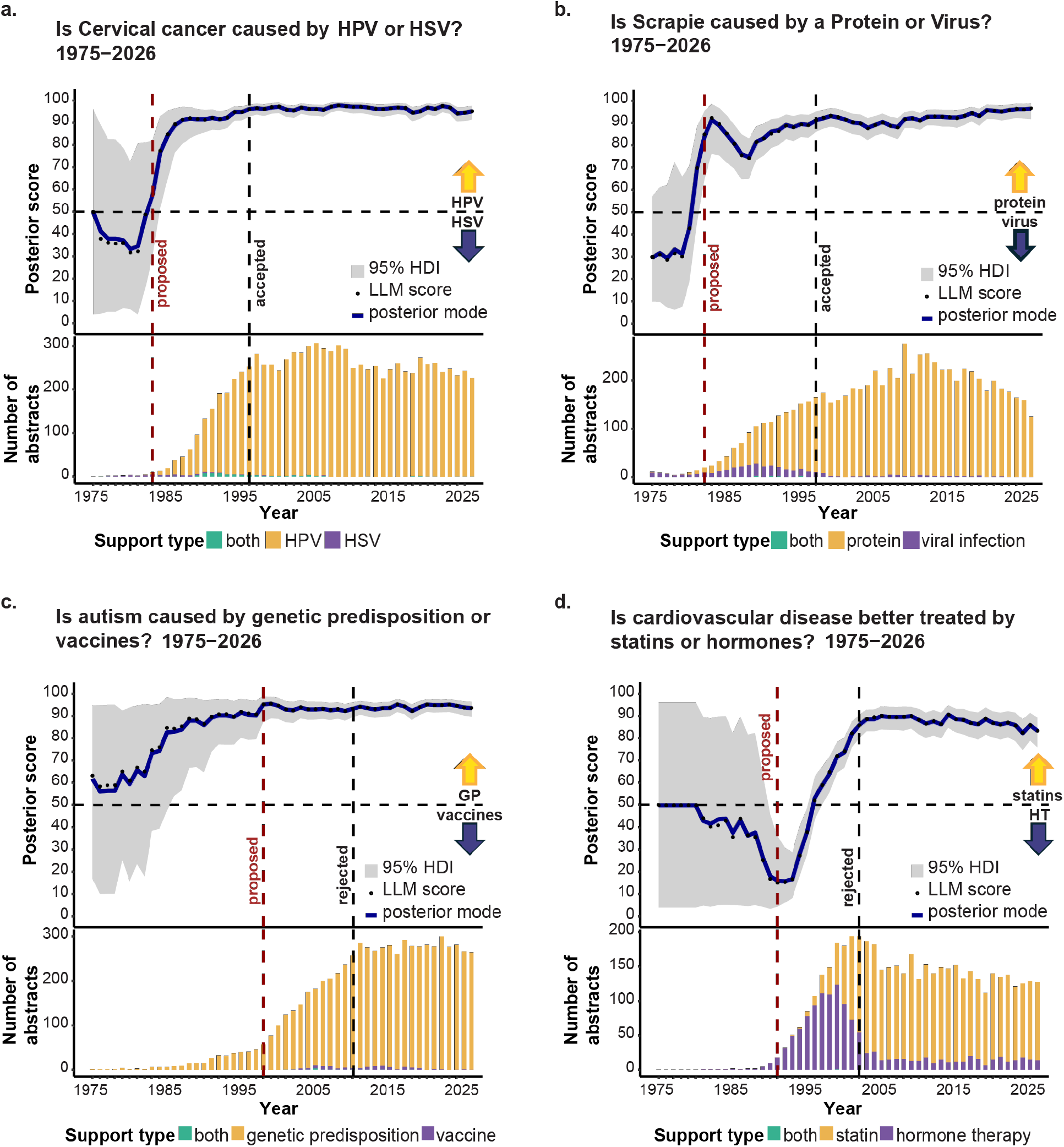
KM-GPT-DCH results on four historically controversial hypotheses. KM-GPT-DCH (iterations=10) comparisons on four historical hypotheses from 1975 to 2026. **For a to d:** The line shows the score posterior mode where a score from 51-100 is more supportive of H1, a score from 0-49 is more supportive of H2, and a score of 50 indicates inconclusive or insufficient information. The gray shaded region is the Bayesian 95% highest density interval of the posterior for each time point (see **Methods**). The bar plot at the bottom shows the number of abstracts used for the evaluation each year with the color scale designating the number supporting the first hypothesis, the second hypothesis, or both hypotheses. Discovery and acceptance dates are based on literature review (see **Supplementary Methods**) **a.** Cervical cancer controversy where the support for the Human papilloma virus (HPV) hypothesis is a score above 50 and support for the Herpes simplex virus (HSV) hypothesis is below 50. **b.** Scrapie controversy where the support for the protein causal hypothesis is a score above 50 and support for the virus causal hypothesis is below 50. **c.** Autism controversy where a score above 50 is more supportive of genetic predisposition (GP) as a cause, and a score below 50 is more supportive of vaccines as a cause. The ‘propose’ x-intercept line indicates when autism was proposed to be caused by vaccines by the 1998 Wakefield paper and the ‘reject’ x-intercept line indicates when the autism/vaccine hypothesis was rejected by the retraction of the Wakefield paper in 2010. **d.** Atherosclerotic cardiovascular disease (ASCVD) controversy where a score above 50 is supportive of statins as a preventive treatment, while a score below 50 is supportive of hormone therapy (HT) as a preventive treatment. The ‘propose’ x-intercept line indicates when HT was proposed to prevent ASCVD, the ‘reject’ x-intercept line indicates when the ASCVD/HT hypothesis was rejected by a large, randomized trial. H1: Hypothesis 1 H2: Hypothesis 2 KM: KinderMiner co-occurrence algorithm DCH: direct comparison of hypotheses GPT: generative pre-trained transformer

Importantly, scores favoring HSV from 1976-1982 indicate that the system’s evaluation is based on the retrieved texts and not its training data (which contains the fact that HPV is the cause of cervical cancer). Thus, not only does the KM-GPT-DCH method support the prevailing hypothesis for a given time but also gives plausible reasoning solely on the time period’s abstracts. In other words, KM-GPT-DCH mitigates parametric memory leakage, as does the SKiM-GPT algorithm on which KM-GPT-DCH is based [36] (see **Supplementary Discussion**).

The third historical set of competing hypotheses involves the causal agent of scrapie (**Fig. 3b)**. Scrapie is a fatal neurodegenerative disease in sheep and goats that takes years to develop. Originally, it was thought to be caused by a slow virus [57], but in 1982 scrapie was proposed to be caused by a protein infection (“prions”) [58]. This proposal was not generally accepted until the mid 1990s [59], with its main proponent Prusiner winning the Nobel Prize in 1997. It is now known that scrapie is caused by a misfolded protein that causes other native versions of the protein to misfold, but this was considered a radical idea at the time as it was thought that causal agents must have nucleic acids [60].

For the potential causes of scrapie, KM-GPT-DCH scores from 1975 to 1979 were generally supportive of viral infection as the leading cause, albeit with very few abstracts. Between 1980-1981 support for protein infection grew, but again with very few abstracts, leading to wide credible intervals. This support for the protein hypothesis occurred before the publication of the landmark paper in 1982 and is based on a study showing that the scrapie “agent” was not inactivated after exposure to UV light wavelengths that should degrade DNA and RNA, thus suggesting that the agent is not a nucleic acid [61]. Despite a slight drop in scores in 1986-1988, from 1982 onwards, scores clearly favored the protein hypothesis (with scores in the 74-97 range indicating strong support for the protein hypothesis), well before the general acceptance date of 1997.

For all three sets of hypothesis pairs discussed so far (for the diseases of peptic ulcer, cervical cancer, and scrapie), KM-GPT-DCH was able to detect the correct hypothesis 3 to 15 years before it was accepted by the scientific community (**Table 1**).

### Evaluating hypotheses that were refuted using KM-GPT-DCH

To show that our algorithm does not just pick the most recent hypothesis or that it can disagree with a hypothesis, we next evaluated two sets of hypotheses that were later refuted. The first example involves the hypothesis that the MMR (measles, mumps, rubella) vaccine caused autism (first published in 1998), which took more than a decade to be fully refuted and retracted in 2010 [47] (**Table 1**). KM-GPT-DCH showed that genetic predisposition was a more likely cause than vaccines for autism from 1988 to 2025 (with pre-1988 having few or no abstracts available, **Fig. 3c**). Additionally, after the year 1988, the mean score per year was above 86, indicating strong support for a genetic cause rather than a vaccine cause. The algorithm never supports vaccines as a cause of autism when evaluating the evidence found in abstracts published during the years of the controversy (1998-2010). When restricting the resolution date to this period or later, the model already resolves genetics as the more strongly supported hypothesis in 1998, 12 years before the retraction of the original vaccine-autism claim. Moreover, when examining earlier resolution dates, genetics has been the most strongly supported hypothesis since 1986, 24 years before the retraction. This conforms to evidence that suggests that autism is a neuropsychiatric condition with one of the highest rates of genetic heritability, estimated between 50 to 85% [62–65].

With the next case we show that KM-GPT-DCH can also choose the more reliable treatment between two hypotheses that change over time. This case involves the use of statins or menopausal hormone replacement therapy (HRT) to reduce atherosclerotic cardiovascular disease (ASCVD) risk. Prior to modern ASCVD reduction therapies (e.g., statins), clinical equipoise existed over whether hormone replacement therapy could reduce ASCVD risk [48]. However, this idea was subsequently refuted in two large randomized clinical trials: beginning with the Heart and Estrogen/progestin Replacement Study (HERS) in 1998 and more comprehensively in the Women’s Health Initiative (WHI) in 2002-2004 [66–69]. Here KM-GPT-DCH reflects the changing knowledge about statin and HRT use for ASCVD prevention. In the early period before the establishment of statins for treatment of LDL-C and reduction of ASCVD risk, HRT was favored as a mechanism to reduce ASCVD risk. However, as statin trials demonstrated clear and beneficial primary and secondary reductions in ASCVD [70–72], and the increased risks of ASCVD events were uncovered for those undergoing HRT [67], statins became dominant (**Fig. 3d**).

In summary, KM-GPT-DCH can correctly refute hypotheses without support (in the case of autism and vaccines) as well as identify temporal trends in therapeutic approaches (in the case of reducing ASCVD risk with the correct identification of statins as the clear preferred treatment as of 1999, three years before the 2002 refutation of HRT as being useful for preventing or reducing ASCVD risk). See **Supplemental Figures** for KM (co-occurrence) only results (**Supplementary Fig. 1**) and KM-GPT non-comparative results (**Supplementary Fig. 2**) for each pair of hypotheses. See **Supplemental Discussion** for more information on each of the above hypotheses, LLM reasoning, and example outputs.

### Evaluating a more recent hypothesis and twenty existing controversial hypothesis pairs with KM-GPT-DCH

A more recent hypothesis about the cause of autism concerns changes in the microbiome, and unlike the vaccine hypothesis, remains unresolved. In the mid 2000s and early 2010s, studies involving mice and microbiota indicated that the gut microbiome can influence changes in behavior in mice [73, 74]. In 2011, a study correlated gastro-intestinal problems and lower levels of certain types of bacteria with autistic children [75]. In addition, short-chain fatty acids, fermentation products of certain bacteria that are associated with autism, were found to induce effects on the brain and alter behavior [76]. Using KM-GPT-DCH, we compared the pre-existing H1 (autism is caused by genetic factors) to the new H2 that autism may be caused by changes in the microbiome. KM-GPT-DCH indicates that from 2013, the score for the microbiome increases (as the genetic predisposition score decreases), but the credible interval stays narrow, and the score never confidently supports the microbiome hypothesis (**Fig. 4a**). As of now, genetic predisposition is still the more plausible causal hypothesis, but the microbiome hypothesis is perhaps worth considering as an additional factor.

**Figure 4.**
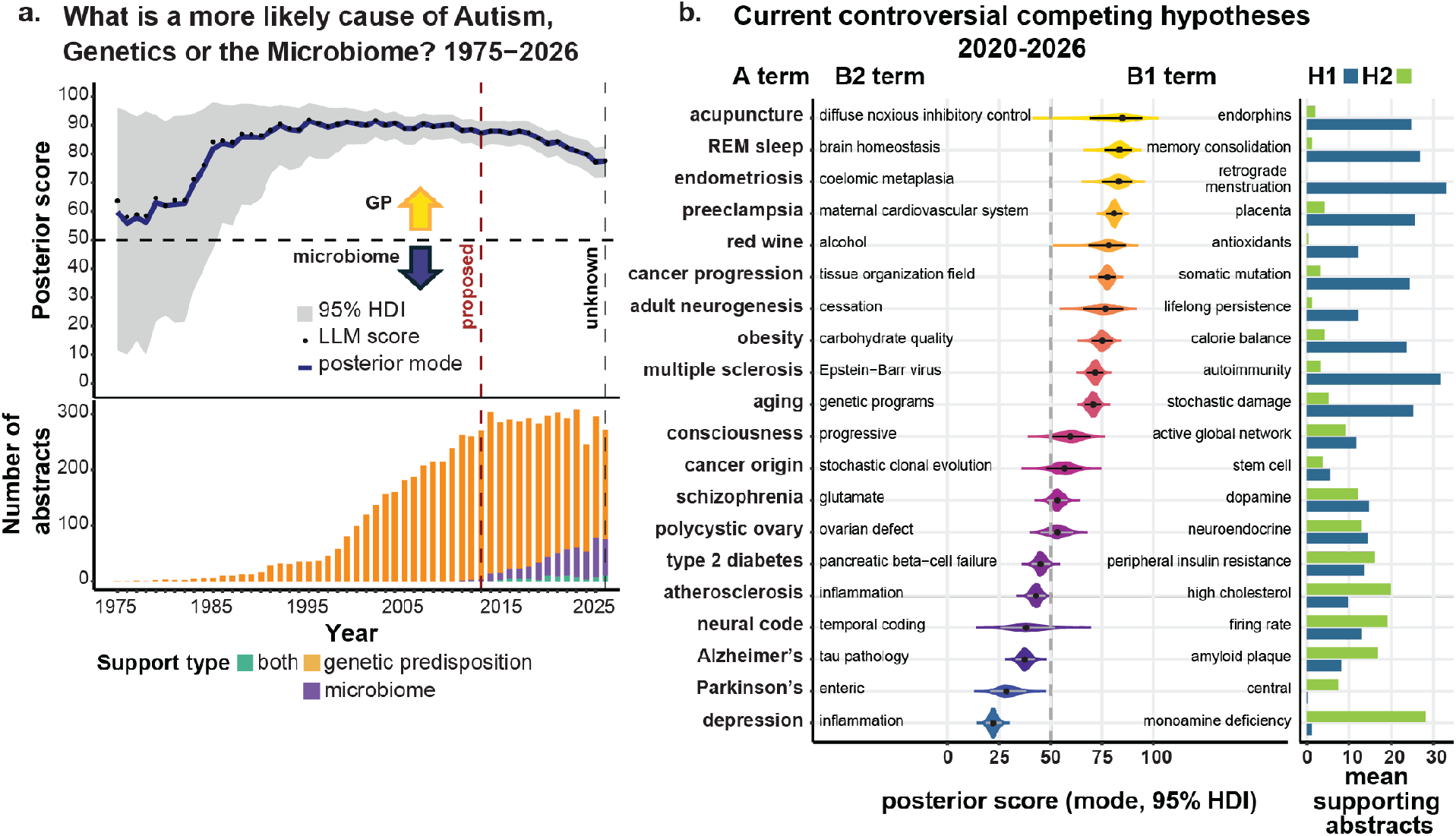
KM-GPT-DCH applied to currently unresolved competing hypotheses. **a.** KM-GPT-DCH (iterations=10) comparisons on the cause of autism, (genetic predisposition (GP) or microbiome), from 1975 to 2026. The graph shows the mode posterior score where a score above 50 is more supportive of H1 (genetic predisposition), a score below 50 is more supportive of H2 (microbiome), and a score of 50 indicates inconclusive or insufficient information. The gray shaded region is the Bayesian 95% highest density interval of the posterior for each time point (see **Methods**). Additionally, the bar plot at the bottom shows the number of abstracts used for the evaluation each year with the color scale designating the number supporting H1, H2, or both hypotheses. Discovery and acceptance dates are based on literature review (see **Supplementary Methods**). **b.** Violin plots of competing hypotheses that are currently unresolved, with literature from the range 2020-2026, where the plots show the KM-GPT-DCH posterior distribution of the scores and the dots in the middle are the mode of posterior scores with error bars representing the 95% credible interval. For each set of hypotheses, scores above 50 are more supportive of H1 and the corresponding B1 term is shown close to the 100 mark, scores below 50 are more supportive of H2 and the corresponding B2 term is shown close to the 0 mark, and a score of around 50 when the credible interval line crosses the 50 mark indicates the competing hypotheses remain unresolved. Bar chart shows mean supporting abstracts from all iterations. H1: Hypothesis 1 H2: Hypothesis 2 REM: Rapid Eye Movement

A primary intended use of KM-GPT-DCH is to help researchers prioritize candidate hypotheses for further investigation by comparing their support in the available literature. To this end, in addition to the autism–genetic predisposition–microbiome hypothesis pair, we evaluated an additional twenty pairs of competing hypotheses that are still undecided using abstracts from 2020 to 2026 (see **Fig. 4b** for all posterior scores and **Supplementary Table 4** for brief descriptions of all hypotheses). Out of these twenty evaluations, six resulted in scores that have credible intervals which overlap with the neutral 50 score line, indicating that neither hypothesis is strongly supported over the other. An example is schizophrenia, a severe mental disorder, where no single cause has been identified. Both dysregulation of dopamine and glutamate signaling pathways have been identified as possible causes [77]. Some studies suggest that both pathways are altered in individuals with the disease [78].

Fourteen of these sets of hypotheses do not overlap with the neutral line, indicating the existing literature leans towards one of the hypotheses. For example, in the case of the underlying biological cause of depression, the competing hypotheses are that it is due to either monoamine deficiency (H1) or neuroinflammation (H2), and KM-GPT-DCH leans heavily towards neuroinflammation with a mode of the posterior score of 22. In this case, while a few abstracts support the monoamine deficiency cause (or both causes), the majority of abstracts sampled (average 96% across 20 iterations) provide support for persistent inflammation as the main cause of anxiety and depression. Many studies have linked pro-inflammatory cytokines such as interleukin-6 (IL-6), with higher levels of depression [79–82]. Additionally, inflammation may induce suppression of monoamines and thus inflammation may be the primary cause [83].

Another set of controversial hypotheses that leans toward one hypothesis is whether neurogenesis occurs into adulthood in the human hippocampus. Prior to the 1960s, neurogenesis was thought to end in childhood, but adult hippocampal neurogenesis gained support in the 1980s and 1990s, first through animal models, then in humans [84]. However, a more recent study from 2018 on post-mortem human brains found no evidence of neurons being generated into adulthood in the hippocampus [85], and since then it has been highly debated. Using our approach targeting papers from 2020 to the present, we found a small number of studies that challenge the view that there is adult neurogenesis. For example, one study found no transcriptomic signatures of neurogenesis in humans compared to other animals, but had a relatively small sample size [86]; another study found that a commonly used marker for newly born neurons, doublecortin (DCX), is expressed in mature neurons throughout the adult brain and does not reliably indicate neurogenesis [87]. Despite this, our algorithm found many studies in support of neurogenesis, including post-mortem immunostaining and single-nucleus RNA sequencing evidence for immature neurons and neural progenitors in adult human brains [88–90]. Overall, our algorithm finds more current papers supportive of adult hippocampal neurogenesis (H1, ∼10 times more), than those papers supportive of cessation of hippocampal neurogenesis in adulthood (H2), and gives a high posterior score of 76.7 in support of lifelong persistence of hippocampal neurogenesis (**Fig. 4b, Supplementary Table 4**).

## Discussion

KM-GPT-DCH extends SKiM-GPT by directly comparing literature support for two user-defined hypotheses and estimating uncertainty in that comparison. We evaluated the system on five hypothesis pairs associated with resolved historical controversies. Across these cases, the system favored the subsequently accepted explanation using literature available before the selected scientific acceptance or rejection milestones (**Table 1, Supplemental Table 1;** see **Supplementary Methods** for the selection of reference dates). These retrospective results illustrate the potential of comparative literature assessment to identify shifts in evidential support before broader acceptance, and suggest the system would accelerate that process in practice.

KM-GPT-DCH provides a general framework for comparing biomedical hypotheses supported by literature in PubMed. The system combines a comparative score with a Bayesian credible interval to characterize relative literature support and uncertainty in that estimate. In our historical evaluations, the tested conversational LLM interfaces and agentic system exhibited citation errors and failures to adhere to publication cutoffs (see **Supplementary Discussion**). In contrast, 100% of the citations in the evaluated KM-GPT-DCH outputs referred to existing PubMed records that adhere to publication cutoffs. By making the retrieved evidence, evaluation procedure, and supporting explanations available for inspection, KM-GPT-DCH provides a transparent and structured basis for comparing hypotheses.

Applied across publication windows, KM-GPT-DCH reveals how relative literature support changes over time. The historical examples include a clear shift toward HPV as a cause of cervical cancer (**Fig. 3a**) and a more nuanced comparison in which bacterial infection was favored as a cause of peptic ulcer while psychological stress remained a contributing factor (**Fig. 2c**). The system consistently favored genetic predisposition over vaccines in the autism comparison during the period of controversy (**Fig. 3c**) and captured the shift toward statins over hormone replacement therapy for cardiovascular risk reduction (**Fig. 3d**). Among twenty unresolved hypothesis pairs, the system favored one of the hypotheses in sixteen comparison pairs, while four pairs remained inconclusive under the analysis criteria. Together, these applications demonstrate the system’s ability to distinguish relative support, characterize unresolved comparisons, and identify changes in the evidence that can inform research priorities.

The web interface (https://skim.morgridge.org/) makes these capabilities accessible to scientists and the public. Researchers can compare specified alternative mechanisms, causal explanations, or interventions, inspect the supporting literature, and use the results to guide further investigation. Comparative scores and uncertainty estimates can help identify where the evidence favors an explanation and where additional studies may be most informative. For scientists exploring unfamiliar domains and members of the public, the interface provides access to the evidence and explanations underlying each comparison. This shared access supports informed examination of biomedical hypotheses and makes the basis of the system’s assessments open to scrutiny. This is an important advantage considering the persistence of controversy among the public even after consensus is built within the scientific community on certain topics – the cause of autism and the potential reasons behind that difference, for example, as addressed in **Supplemental Discussion**.

The KM-GPT-DCH algorithm has some limitations. First, the KM, KM-GPT, and KM-GPT-DCH algorithms do not currently search the main body of the papers, tables, figures, supplemental materials, or any numerical data. We plan to include some of these other data sources in subsequent updates. Second, when both hypotheses have little literature support, evaluation may be difficult because of low statistical power.

Third, the algorithm can only compare two hypotheses at a time. An example of when comparing three or more hypotheses would be useful is the case of peptic ulcer. In the 1990s, it became clear that NSAIDs (non-steroid anti-inflammatory drugs) also can be a primary cause of peptic ulcer [91, 92], and these ulcers can be exacerbated by psychological stress [93]. Thus, there are three hypothetical causes for peptic ulcer: psychological stress, bacterial infection, and NSAIDs. Finally, the A terms and B terms used to search for a given hypothesis can impact the results; see **Supplementary Methods** for guidance on choosing terms.

## Conclusions

KM-GPT-DCH expands the capabilities of SKiM-GPT from evaluating individual hypotheses to directly comparing their relative literature support, with Bayesian estimates that quantify uncertainty in the comparison. Together, these capabilities help researchers distinguish evidence favoring one hypothesis from comparisons that remain inconclusive. Applied across publication windows, the system tracks changes in relative support and the uncertainty surrounding those changes. Retrospective analyses demonstrate its potential to identify support for subsequently accepted explanations before selected historical acceptance milestones, while applications to unresolved hypothesis pairs illustrate its use in prioritizing further investigation. KM-GPT-DCH therefore provides a structured, inspectable framework for comparing competing hypotheses and informing research priorities.

## Declarations

### Ethics approval and consent to participate

Not applicable

### Consent for publication

Not applicable

### Availability of data and materials

All code is available through GitHub: https://github.com/stewart-lab/skimgpt.

**Supplementary Data 1**: ***Detailed Q and A web-based LLMs.*** All questions and answers posed to and given by the chat-based LLMs and Robin for evaluation that are summarized in Supplementary Tables 2 and 3.

**Supplementary Data 2: *All KM-GPT-DCH outputs.*** Includes all responses for every time point and iteration from KM-GPT-DCH for each set of hypotheses discussed, can be found: Zenodo DOI: **10.5281/zenodo.20547913.**

**Supplementary Data 3**: All results from the comparative hypotheses run for variance testing.

### Author Contributions

Ron Stewart conceived the work. Bethany M. Moore, Jack Freeman, Ron Stewart, and Robert J. Millikin helped to design the work. The acquisition, analysis, and interpretation of data was done by Bethany M. Moore, Jack Freeman, Robert J. Millikin, Chitrasen Mohanty, Kevin Shine George, Aviral Bal, John-Demian Sauer, Megan E. Spurgeon, Darcie L. Moore, Brittany G. Travers, and Ron Stewart. The software for this project was created by Bethany M. Moore, Jack Freeman, Robert J. Millikin, Chitrasen Mohanty, Kevin Shine George, Aviral Bal, Cannon Lock, and Ron Stewart. The study was written and revised by Bethany M. Moore, Jack Freeman, Robert J. Millikin, Chitrasen Mohanty, John-Demian Sauer, Megan E. Spurgeon, Darcie L. Moore, Brittany G. Travers, and Ron Stewart.

## Acknowledgements

We thank Debbie McKenzie (University of Alberta) for insightful comments on the history of the prion hypothesis. We thank Alicia Williams for assistance with manuscript editing.

