## Supplementary Methods for "Quantifying Support for Competing Biomedical Hypotheses using Large Language Models and Bayesian Analysis"

### Supplementary Discussion

#### What is temporal filtering and why do we need it?

Temporal filtering is a popular method for evaluating literature-based discovery (LBD) systems [1]. In temporal filtering, the LBD system is restricted to only have access to papers or abstracts up to a particular cutoff date, which is typically before the discovery is made. The LBD system is considered to have been successful if it is able to suggest a particular discovery (such as a drug would be useful for treating a disease) using literature from before the discovery is made. While we are not performing LBD here, we use temporal filtering to allow KM-GPT-DCH to evaluate hypotheses only using literature at or before a particular date. Importantly, co-occurrence model-based algorithms like SKiM [2] can perform temporal filtering easily by only allowing papers published before a particular date to be included in the analysis. However, LLMs out of the box are not well-suited for temporal filtering because of leakage from parametric memory (see next section). Parametric memory is based on the training data. The training data and thus the parametric memory is typically not time stamped.

#### What is parametric memory leakage?

Parametric memory leakage (leakage), in the context of KM-GPT-DCH, refers to the unintended use of information stored in the parameters and weights of a large language model (LLM) to generate results without retrieval-augmented generation (RAG). This is also called parametric memory [3]. This occurs because LLMs are trained on large, untimed datasets, making them unsuitable for temporal analysis when not constrained by RAG. LLMs, without external frameworks, cannot effectively be used for temporal filtering analysis as the weights and parameters of the LLM are not time stamped. For instance, we can ask an LLM (with no RAG) Does HPV cause cervical cancer? We would expect the LLM to answer in the affirmative because frontier LLM’s training corpus contains documents published/written after HPV was discovered as the primary cause of cervical cancer. Proving the absence of leakage when RAG is applied is crucial to ensure KM-GPT-DCH’s viability and accuracy, as it guarantees that hypothesis evaluations rely exclusively on the provided, contextually relevant literature, rather than pre-existing biases or untimed knowledge within the model. We have previously shown that our RAG and structured output approach avoids parametric leakage [4].

Additionally, using our time filtered approach, we can show how our model avoids parametric leakage. In the subsection of cervical cancer below, we discuss how from years 1975-1982, the model supports HSV as the main cause of cervical cancer, even though it knows that HPV is the actual cause in its parametric memory. Instead of using the LLM knowledge itself, it is paying attention to the abstracts in front of it. Other examples from our main hypothesis comparisons run also exist. In the case of peptic ulcer, in 1980 it gets a score of 30, favoring stress. If there were parametric leakage it would favor bacterial infection because as of 2026, this is a known cause while stress is only a contributing factor. The following score rationale highlights the LLM’s reasoning in this case: “Two abstracts (PMID: 7432255, 7388642) explicitly cite psychological stress/personality conflicts as facilitating or predisposing to duodenal ulcer, supporting H2. No abstract directly attributes peptic ulcer causation to bacterial infection; one (7425593) merely reports bacteria present, without causal claim classified as inconclusive. Following guidelines, with moderate support for H2 and none for H1, assign a score slightly below the 25 anchor (H2 three-times more likely) to reflect modest dataset size.” The final example of avoiding parametric leakage is with the scrapie competing hypothesis example. Our model gives the scrapie hypotheses a score of 15 in favor of a virus cause in 1977 before the protein hypothesis existed, with the following rationale: “Only abstract 404706 repeatedly characterizes scrapie as a slow-virus disease and details virus particles and their replication, directly supporting Hypothesis 2 and contradicting Hypothesis 1. No abstract offers protein-only causation evidence. With unilateral, clear support for H2, the evidence strongly favors H2; limited dataset tempers maximal certainty.” These examples collectively show that our system can avoid directly relying on the weights and parameters of the LLM to answer a question by employing RAG instead, an approach that is important because it enables effective temporal filtering [5], as demonstrated in our temporal filtering analysis.

#### Quantifying Uncertainty in LLM scores

Quantifying uncertainty of an answer, a confidence interval or *p*-value of that score, by a frontier LLM is challenging. Existing methods to assess the uncertainty of an LLM fall mainly into the categories of verbalization, sequence probability, surrogate token, consistency, and trained probing [6]. Verbalization relies on directly asking the LLM the level of confidence it has in its response. This method is fairly accurate, even surpassing sequence probability methods [7, 8]. Sequence probability methods use the log probabilities from the output tokens to estimate confidence [9]. A drawback to this method is that the confidence predicted can be about how the statement is made, not about the actual statement [6]. Additionally, frontier LLMs no longer expose their log probabilities of output tokens, which are also needed for surrogate token methods and consistency methods. Surrogate token methods involve combining verbalization and sequence probability by finding the probability of one output token, like the probability of the output token to the proposition, ‘is this correct?’ [10]. Consistency methods involve sampling multiple answers to a question of statement and measuring the consistency between answers [11]. However, these methods do not provide a confidence score, it just gives the most consistent answer. Trained probing methods employ machine learning to extract features from hidden layers of the LLM and then train a classifier on these features to determine if the statement is true or false [12]. However, in a frontier LLM, the hidden layers are not exposed and therefore cannot be used to train. If hidden layers can be used, such as in open source models, it is often difficult to determine which layer(s) to use [13]. This type of method also requires curated training data.

Recently, researchers have employed Bayesian methods to assess the credibility and stochasticity of LLM responses as Bayesian methods have advantages over traditional frequentist methods such as assessing probabilities of a response, and have shown some promise in quantifying LLM output uncertainty [14, 15]. We therefore turn to Bayesian inference for our estimation of uncertainty in our system. See the main text for details about our Bayesian method.

#### Conversational LLM interfaces for evaluating competing hypotheses

To see differences between our method and asking an LLM via a conversational interface, we tested four different commonly used: ChatGPT(gpt-5.0)-5 and ChatGPT-4o, Perplexity (Sonar (built on Meta Llama 3.3 70B) with routing from Mistral Large 2 and Gemini 2.5 Flash), Gemini (2.5 Flash), and DeepSeek (R1-0528), and one multi-agentic system: Robin, using the Crow interface (<https://platform.futurehouse.org/>). To determine how well each LLM performed, we checked the accuracy of the citations provided and whether the LLM could determine the correct hypothesis at a given time. When attempting to ask the LLM via chat, we found there were consistently four types of errors: 1) the reference would be after the cut-off date, 2) the citation was incorrect or incomplete, 3) the reference was non-existent, or 4) the reference was not relevant. To test, we used two cut-off dates, the first being two years before the landmark study was published (prior), the second being three years after the publication came out (post). For example, the peptic ulcer example (landmark paper in 1984) has a prior cut-off date of 1982, and a post cut-off date of 1987. The prior cutoff date should represent the previous hypothesis (before evidence for the new hypothesis) and the post cutoff date should represent the new hypothesis (after evidence for the new hypothesis is published). The overall response result and reference evaluation for the LLM-interfaces is summarized in **Supplementary Table 2,** and all LLM answers are found in **Supplementary Data 1: *Detailed Q and A Web LLMs***.

For the first type of error, we found LLMs via chat interface could not correctly perform temporal filtering and would consistently reference websites, blogs, and papers after the cut-off date to come up with an answer. We found that 60.4% of the total number of prior references fell into this error category, and in particular, the references from Perplexity tended to be in this category, where 39 out of 41 cases used references beyond the cutoff date. For the post-study cut-off date, 51.0% of references fell into this category. This highlights that using an LLM via chat interface may be insufficient for comparing current competing hypotheses. Interestingly, this happens even when using a post-study cut-off date. Many sources in this category are summaries of the findings like Wikipedia entries, websites, or reviews, indicating that the LLM uses summarized information frequently after the cut-off date to choose the correct hypothesis.

In the second error type, reference citations were often incorrect, with incorrect authors, titles, or DOI URLs. This occurred with 11.3% of the references cited by LLM chat interfaces for the prior study cut-off date and 16.0% of the references cited for the post study cut-off date.

In the third error type, references are completely fabricated. 11.6% of the references prior to the landmark study and 5.6% after publication were completely fabricated and did not exist at all.

Finally, the fourth error type represents irrelevant citations. 1.7% (prior) and 1.5% (post) of the references cited by the LLMs were not relevant to either of the hypotheses being tested.

In total, only 15.0% of references for the pre-study cut-off date and 25.9% for the post-study cut-off date were cited correctly and were relevant. We found that overall citation accuracy was better for Robin (tested only on the cervical cancer hypothesis), although not completely without error (~85% total legitimate references for both prior and post cut offs, **Supplementary Table 3**).

Additionally LLMs by themselves cannot always get the correct answer, as they tend to use parametric data, cannot time-filter their answer (using parametric information stored in the model based on training documents after the cut-off date), and again may hallucinate if they do not know the correct answer [16]. When testing LLM-interfaces on correctness using all hypotheses, before the landmark study came out, the LLMs only provided the correct response (the pre-existing hypothesis) 66.7% of the time. Even after the published study came out (but before general acceptance, the LLMs found the correct response (the new hypothesis) only 80% of the time (**Supplementary Table 2)**. Additionally, Robin tended to choose the new hypothesis before it had been proposed (50% of the time, **Supplementary Table 3**). This shows that these systems likely used data already present within the trained model to answer the question, thus making it unsuitable for historical analysis or for analysis where it is critical to restrict to only trusted sources of information (such as PubMed). These analyses demonstrate an inability to perform temporal filtering and the common hallucinatory problem with LLMs [17, 18] that hinders the model’s interpretability and reliability.

Finally, both LLM chat systems and agentic systems tended to cite a narrow breadth of evaluated scientific papers (~12-13 on average for LLM chat systems (**Supplementary Table 2**) and ~8 for Robin (**Supplementary Table 3**)). This contrasts with KM-GPT-DCH, which can obtain a wider breadth of papers (50 abstracts at one iteration for this study, and with 10 iterations with abstracts sampled with replacement, the algorithm can evaluate hundreds of abstracts). KM-GPT-DCH also has a finer resolution scoring system and a credible interval. See **Supplementary Data 1**: ***Detailed Q and A web-based LLMs*** for detailed questions and responses from web-based LLMs and agentic systems in evaluating competing hypotheses.

There are several advantages to the KM-GPT-DCH system over chat-based LLM systems:

1. No hallucinated citations and greatly reduced LLM hallucinations in reasoning and evaluation.

2. For KM-GPT-DCH, the exact text that the LLM is reasoning about is transparent, which is not necessarily true for other LLM-based implementations.

3.  Parametric memory leakage is not an issue with KM-GPT-DCH but is a primary source of information for other LLM implementations. This makes determining the sources contributing to the output challenging and temporal filtering impossible.

4. Importantly, KM-GPT-DCH tightly controls dates and allows true temporal filtering to understand the history of a hypothesis and to be able to accurately assess the support for a hypothesis at a particular time.

5. Chat-based LLM implementations choose the papers to discuss, and it isn’t clear how they are chosen. In KM-GPT-DCH, the system reliably chooses the articles to reason about based on co-occurrence and temporal filtering dates.

6. KM-GPT-DCH can be as comprehensive as the user desires. It can evaluate 5 abstracts, or 500 abstracts. It is not generally possible to tune the level of comprehensiveness in web-based systems.

#### Other systems for hypothesis evaluation

Other systems use retrieval-augmented generation (RAG) to use published literature to aid in hypothesis evaluation. These include OpenScholar [19], ValSci [20], and proprietary systems such as Scite [21], Elicit (<https://elicit.com/>), and Consensus (<https://consensus.app/>). Tools like ValSci and OpenScholar focus primarily on claim verification, where LLMs query a database or search for papers that may support or refute a single statement. The proprietary Consensus tool uses a "Consensus Meter" to provide an aggregate distribution of supporting versus contesting evidence, yet this remains a statistical summary of search results rather than a comparison of two distinct theories. Similarly, Elicit excels at automated data extraction through organizing study characteristics into tables to assist in systematic reviews, but it leaves the higher-level task of weighing competing hypotheses to the human researcher. Scite offers citation context by identifying whether a paper is supported or contrasted, but it operates at the metadata level, lacking a framework to evaluate the underlying scientific validity of one hypothesis relative to another. As mentioned in the main text, reliance on evaluating hypotheses in isolation tends to lead to score shrinkage (where individual evaluations tend to get similar scores) and thus the inability to effectively compare cases [22–24]. Thus, the advantages of KM-GPT-DCH are that it can effectively compare hypotheses as well as give an estimate of confidence to its evaluations.

#### KM co-occurrence results

Below are results from the KM model on the other four main historical hypothesis pairs (other than peptic ulcer which was mentioned in the main text). In many of the cases, KM results in the correct term being favored at a given time but of course cannot understand the relationship of the two terms.

Cervical cancer

With the co-occurrence model (KM), we use the two-proportion z-test (z-interval) to determine if there is a difference between the proportion of abstracts relevant to the new hypothesis (HPV) and the proportion of abstracts relevant to the preexisting hypothesis (herpes simplex virus or HSV). During this time (1983-1996), we see with the co-occurrence model (**Supplementary Fig. 1a**) that more papers were published with cervical cancer associations with HPV than HSV, and the proportion is significantly higher by 1984 (*p* <0.05). There was an even stronger association of HPV with cervical cancer when compared to HSV by 1987 (*p* < 1.8e-52), well before the time of general scientific acceptance of HPV as causal to cervical cancer in 1996.

Scrapie

Similarly, we used KM to look at the second pair of competing hypotheses of whether scrapie is caused by a protein or a virus infection. We found that KM could detect a significantly higher proportion of abstracts associating scrapie with protein infection than with viral infection by 1992 (*p* < 0.05). By 1996, co-occurrence with protein infection was very strong (*p* < 1.5e-5) (**Supplementary Fig. 1b**), indicating that before the general scientific acceptance date of 1997, protein infection was significantly more highly associated with scrapie than viral infection.

Autism

KM shows strong co-occurrence numbers with autism and vaccine link in 2002 and for some years after with the ratio favoring vaccines until 2007, although this ratio is never significant at $a=0.05$. In this case, some of these papers in which vaccines and autism co-occur after 1998 address and refute the claim that vaccines cause autism, causing the co-occurrence ratio to increase even though the actual support of this claim decreases (**Supplementary Fig. 1c)**. In 2008 and after the z-score remains positive through current time (favoring the genetics term), indicating before the Wakefield paper was retracted in 2010 there was already significantly higher associations with autism and genetics than autism and vaccines. We also see a drop-off of ‘autism and vaccines’ papers to zero starting in 2014. This is because at this point, Fisher’s Exact Test no longer finds a significant number of ‘autism and vaccines’ abstracts relative to overall vaccines abstracts (*p*-value < 0.05), and we do not obtain any counts that are past this cutoff. This highlights a downside to using *p*-value cutoffs as they can distort the data based on what is deemed significant. It is also another reason KM-GPT-DCH is a better approach as this method employs a Bayesian credible interval around a score, but not a hard cutoff of when a result is considered significant or not.

ASCVD (Atherosclerotic Cardiovascular Disease)

Finally, KM analysis compares HRT (hormone replacement therapy) with statins showing statins with a significantly higher abstract count than hormone therapy by 1993. This trend continues to climb in favor of statins through 2026 (**Supplementary Fig. 1d**). This example underscores the importance of temporal trends in therapeutics. Prior to statin use for ASCVD risk reduction, clinical equipoise existed as to whether HRT could be used as a method to reduce ASCVD risk, but following the results of randomized controlled trials, HRT was no longer viewed as a method for ASCVD risk modification.

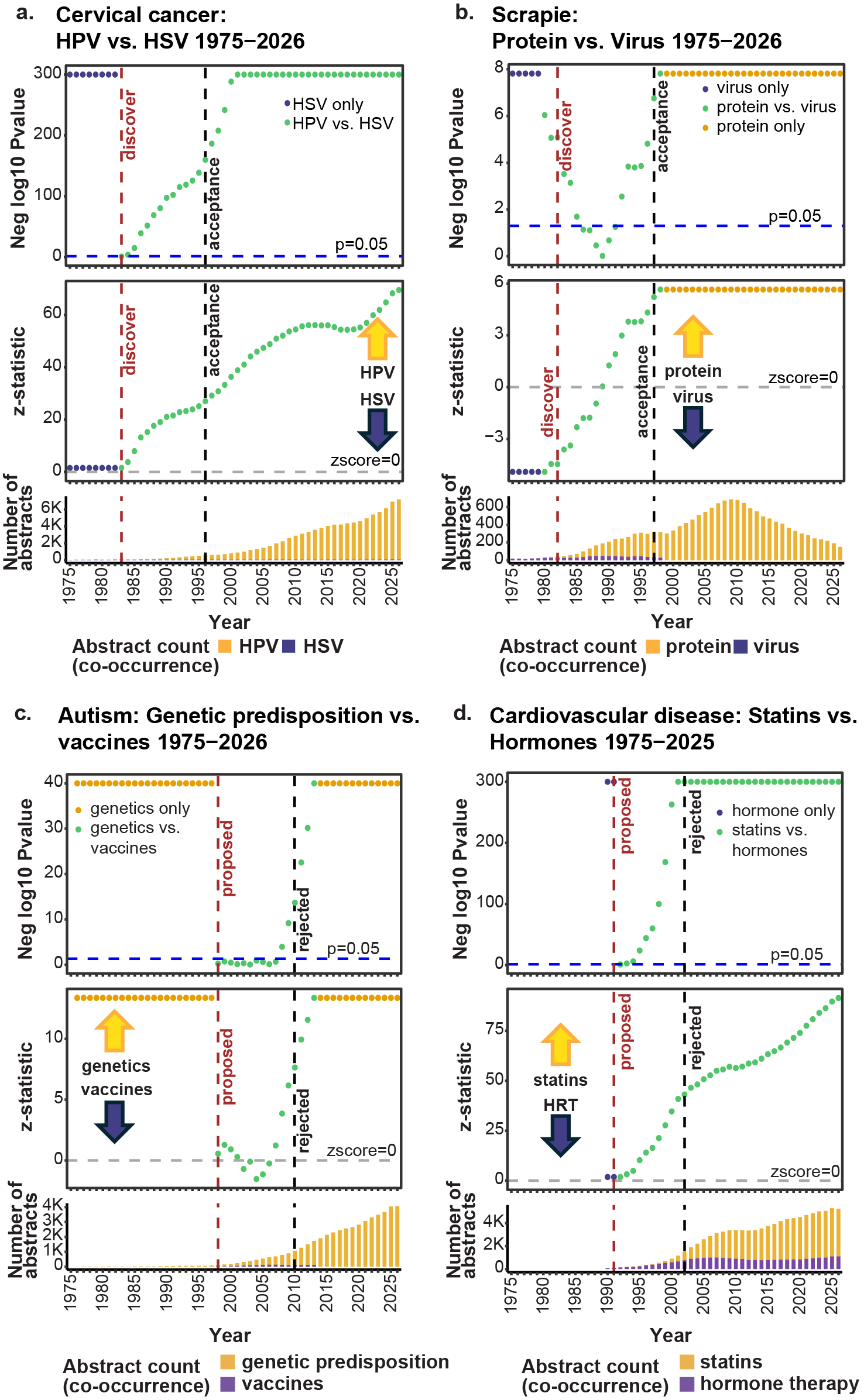

**Supplementary Figure 1**: **KinderMiner with ratio of ratios z-statistic**

All panels (**a,b,c,d**) show the results from the two-proportion *z*-test for each year between the sets of competing hypotheses (H1 vs. H2) using the KM algorithm from 1975 to 2026, run cumulatively. The top panels show the negative log10 of the *p*-value, where the dashed y-intercept line is the adjusted *p*-value of 0.05. The middle panels show the *z*-statistic, where the dashed line represents *z*-statistic of 0, where above zero the algorithm favors H1 and below zero the algorithm favors H2. Finally, the bottom panels show the number of abstracts obtained for each B-term that co-occur with the corresponding A term.

**a**. KM results from cervical cancer causation hypotheses; H1: Human Papillomavirus (HPV), H2: Herpes Simplex virus (HSV). The ‘discover’ x-intercept line indicates when HPV was found as a causal agent and the ‘acceptance' x-intercept line indicates when the HPV cause was accepted by the scientific community. **b.** KM results from scrapie infection causation hypotheses; H1: protein infection, H2: virus infection. The ‘discover’ x-intercept line indicates when proteins were discovered to cause infections like scrapie and the ‘acceptance' x-intercept line indicates when prions were accepted as a causal agent. **c.** KM results of autism causation hypotheses, where genetic predisposition is H1 and vaccines is H2. The ‘propose’ x-intercept line indicates when autism was proposed to be caused by vaccines by the 1998 Wakefield paper and the ‘reject’ x-intercept line indicates when the autism/vaccine hypothesis was rejected by the 2011 retraction of the 1998 paper. **d.** KM results of atherosclerotic cardiovascular disease (ASCVD) preventative treatment hypotheses, where statins are H1 and hormone replacement therapy (HRT) is H2. The ‘propose’ x-intercept line indicates when hormone therapy was proposed to reduce ASCVD risk, the ‘reject’ x-intercept line indicates when the ASCVD/hormone therapy hypothesis was rejected by a large, randomized trial.

H1: Hypothesis 1

H2: Hypothesis 2

KM: KinderMiner co-occurrence algorithm

—------------------ end of figure legend —------------------

#### Running KM-GPT

When KM-GPT is utilized, positive results for both hypotheses are usually produced. This is largely because when KM obtains abstracts for only one of the hypotheses, there may be correlative or weak evidence for the hypothesis and is not directly refuted, thus deserving a perhaps lower but still positive score.

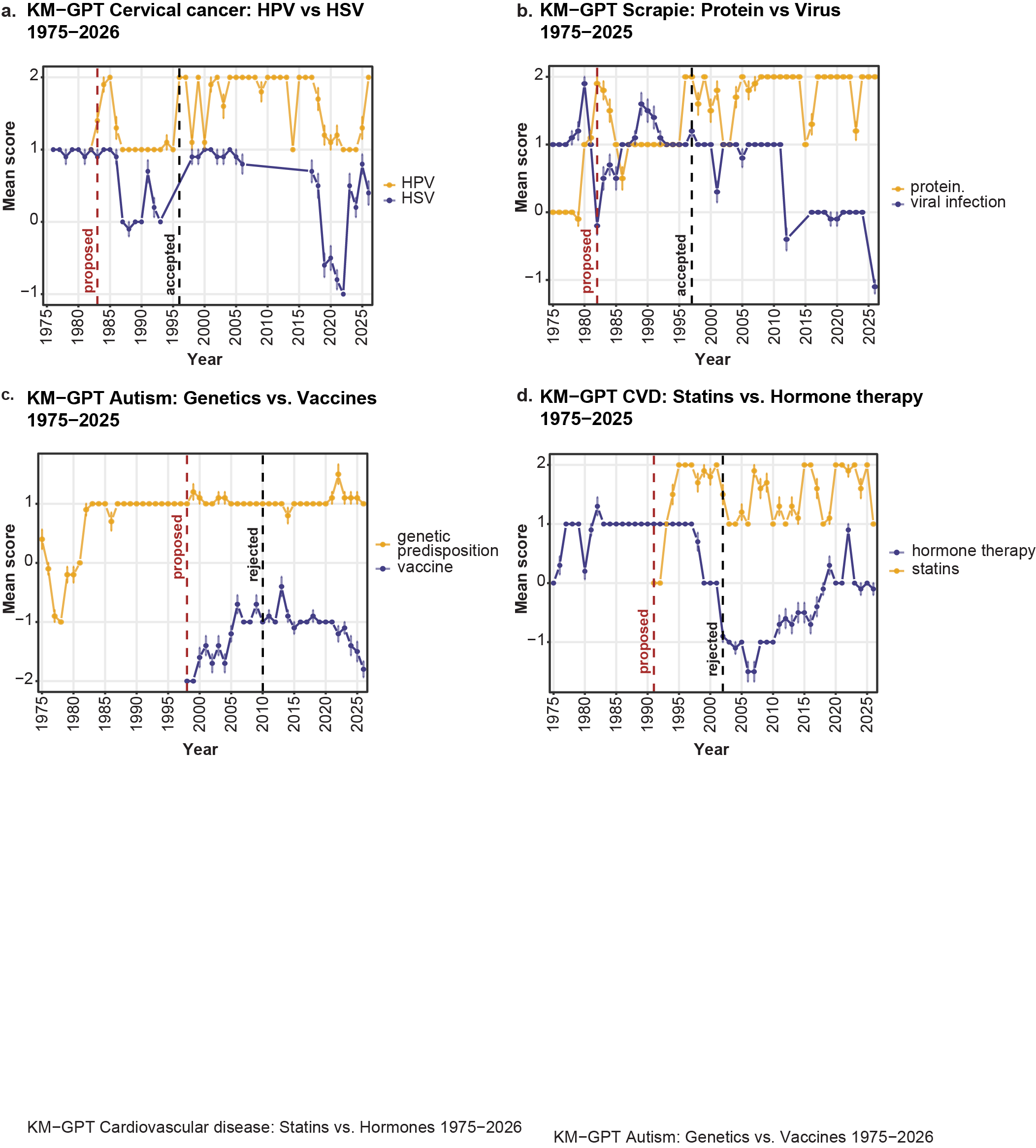

**Supplementary Figure 2. KM-GPT runs**

KM-GPT scores over time (1975-2026) for each hypothesis when run as a single hypothesis, run with ten iterations, where each point is the mean score and error bars show standard error across iterations. The y-axis shows the score value ranging from +2 to -2, where +2 KM-GPT strongly agrees with the hypothesis and -2 KM-GPT strongly disagrees with the hypothesis. The x-axis shows the given year, and the lines are color coded for one of the hypotheses. **a.** The cause of cervical cancer, HPV (yellow) or HSV (purple). **b.** The cause of scrapie, protein (yellow) or viral infection (purple). **c.** The cause of autism, genetic predisposition (purple) or vaccines (yellow). **d.** The best treatment for cardiovascular disease, statins (yellow) or hormone therapy (purple). Note that after the mid 2000s, HRT was no longer viewed as a treatment for reducing ASCVD risk.

KM-GPT: KinderMiner with a Generative Pretrained Transformer.

—------------------ end of figure legend —------------------

In the case of cervical cancer, when running KM-GPT, positive scores are produced (with a range from -2 to +2, where +2 is highly supportive of the hypothesis and -2 indicates the hypothesis is strongly refuted) for both HSV and HPV, including positive scores for HSV as a causal agent well after general acceptance that HPV is the most likely cause of cervical cancer (**Supplementary Fig. 2a**). This is because HSV, while not the causative agent, is still strongly correlated with cervical cancer. If the LLM is not comparing this cause to anything else and only has abstracts citing correlations of HSV and cervical cancer, it will give this a positive score for being a cause. Thus KM-GPT-DCH provides a better evaluation of the cause of cervical cancer, as KM-GPT-DCH evaluates abstracts for both HSV and HPV at the same time, and can choose the more likely hypothesis.

Similarly, when using KM-GPT to evaluate the cause of scrapie, positive scores are produced for both protein and virus as causal agents, including some positive scores (although also some negative scores) after the acceptance date (**Supplementary Fig. 2b**). When looking at the abstracts from 2007 when the viral hypothesis gets a high score, there are indeed abstracts that support a viral hypothesis (1-33 abstracts), but more usually support a protein hypothesis (14-34 abstracts). Therefore, if run in a non-comparative way, the LLM only sees abstracts relevant to one of the hypotheses, which would make one hypothesis seem plausible even if the other hypothesis has more evidence. When running KM-GPT on the autism causal hypotheses of genetic predisposition or vaccines, the vaccine hypothesis never gets a positive score while genetic predisposition gets a high positive score (+2) starting from the early 1990s continuing through today (**Supplementary Fig. 2c**). This is a case where KM-GPT is sufficient and indicates that there is little to no evidence that vaccines cause autism. Thus, when we compare two hypotheses, KM-GPT-DCH can choose the overall better one, while KM-GPT is not always able to clearly identify the better hypothesis. See **Supplemental Data 2** for all KM-GPT runs.

#### LLM reasoning for hypotheses

For each hypothesis, the LLM provides reasoning along with the score. Here the reasoning and the references used for one historical hypothesis pair is provided, to show that 1) the reasoning is indeed sound and 2) that the LLM bases its decision on the abstracts of the given time and not its parametric data.

##### Cervical cancer

As mentioned in the main text, HSV was the dominant theoretical cause of cervical cancer from 1975-1982, for which our algorithm scored in favor of HSV in these years. In the years following the report by Durst *et al.* in 1983 [25] HPV was hypothesized as the leading cause, during which KM-GPT-DCH scored in favor of HPV. From 1976 to 1982, the mean score of the posterior modes (37) favors HSV and in 1982, two to four abstracts per iteration support the HSV cause while only one abstract has support for HPV. The reasoning from 1982 states that one study showed there were higher amounts of HSV-2 antibodies in cervical cancer patients compared to matched controls [26] while another study showed that HSV-2 is linked to an increased risk in cervical cancer [27]. In 1984, when the algorithm favored HPV (mean score of 77.5), there were eight abstracts that support HPV as the cause of cervical cancer while only three to four support HSV. Some of the abstracts supporting HPV provided molecular evidence, including the landmark paper that isolated HPV DNA from cervical cancer biopsies [28, 29] while the abstracts supporting HSV-2 only show correlative antibody associations with cervical cancer or support via a mouse model [27, 30, 31]. The reasoning confirms that the algorithm is basing its decision on these abstracts and the correct time period with no parametric memory leakage. See below for one example of results from a 1984 run by the KM-GPT-DCH algorithm and **Supplementary** **Table 1** for a list of the various hypotheses and when/if they were resolved by KM and KM-GPT-DCH. For all KM-GPT-DCH outputs, see **Supplemental Data 2**.

##### Peptic Ulcer

The peptic ulcer causation hypotheses are not as straightforward because, as mentioned in the main text, while *H. pylori* is the bacterium that is the main causal agent to ulcers, psychological stress is also an aggravating factor. KM-GPT-DCH results vary, but eventually by 1987 KM-GPT-DCH consistently favored the bacterial hypothesis as the better one. As shown in the result example below, in 1987 multiple studies found *H. pylori* in duodenal ulcers [32–35], and while some studies found correlation with psychological stress [36], others are inconclusive or contradictory [37, 38]. The algorithm correctly evaluates that the bacterial hypothesis was stronger with evidence from multiple studies, whereas evidence for a psychological stress causal component was weaker and, in some studies, refuted.

Additionally, on average the proportion of abstracts favoring one hypothesis over another mirrors the LLM score and posterior score (see timecourse_data.csv tables in KM-GPT-DCH output folders in **Supplemental Data 2**), however in certain instances the LLM score better aligns with the content and quality of the abstracts, not just the quantity or proportion. For example, in the peptic ulcer run of 1987, the mean LLM score is 63 and the posterior score is 62.9 favoring the bacterial infection hypothesis (H1), but the proportions favor psychological stress (H2) with on average 7.5 abstracts supporting H1 and 11.8 abstracts supporting H2 (a proportion of 61% in favor of H2). However, as we can see in the reasoning example below in the **Example KM-GPT-DCH results** section for peptic ulcer, the reasoning for the 1987 run states “direct, disease-localized bacterial findings modestly outweigh the larger but less causally specific and mixed psychological literature”. Therefore, the reasoning can distinguish “better” support from simply more abstracts supporting a given hypothesis, and we tend to agree with the overall reasoning of the LLM given the abstracts it evaluated.

#### Example KM-GPT-DCH results

##### Cervical cancer

The following is an example of KM-GPT-DCH results for a 1984 run of the cause of cervical cancer, hypothesis 1: cervical cancer is caused by HPV, hypothesis 2: cervical cancer is caused by HSV, iteration 8. 1984 is a year after the landmark paper came out showing HPV as a causal agent, and the time when the HPV hypothesis clearly becomes the favorable hypothesis.

Result per abstract:

- PubMed ID: 6399280
  - evidence: Herpes simplex virus type 2 remains a candidate oncogenic agent, but comparative seroepidemiological results among cases and controls are inconsistent. Human papillomaviruses are clearly more important.
  - label: supports_H1
- PubMed ID: 6304740
  - evidence: HPV 16 DNA was detected under high-stringency conditions in 61.1% (11/18) of German cervical-cancer samples and 34.8% (8/23) of samples from Kenya and Brazil. Only 2 of 33 benign condylomata contained HPV 16 DNA; the data indicate HPV 16 DNA prevails in malignant tumors.
  - label: supports_H1
- PubMed ID: 6318956
  - evidence: The gradual transition and morphologic overlap between subclinical papillomaviral infection and CIN support the postulate that they are a single disease spectrum. Inverse relationships between late viral expression and nuclear distortion fit experimental models of viral oncogenesis.
  - label: supports_H1
- PubMed ID: 6305251
  - evidence: Virus was found in 83% of pure condylomas, 16% of dysplastic condylomas, and 0% of condylomatous dysplasias. The viral absence with dysplastic features could be a new argument to suspect the carcinogenic potential of HPV for the cervix and external genitalia.
  - label: supports_H1
- PubMed ID: 6329740
  - evidence: HPV 18-related DNA was detected in 9/36 cervical carcinomas from Africa and Brazil and 2/13 cervical tumors from Germany, while benign tumors, carcinoma in situ, and normal cervical biopsies lacked detectable HPV 18 DNA. In cervical-cancer biopsies, HPV 18 DNA might be integrated into the host cell genome.
  - label: supports_H1
- PubMed ID: 6100284
  - evidence: The abstract reports an association of HPV-6, -10, -11, -16, and -18 with cancers of skin and mucous membranes, including cervical cancer.
  - label: supports_H1
- PubMed ID: 6299907
  - evidence: Subclinical HPV infection and cervical cancer have essentially identical epidemiological characteristics, and SPI commonly coexists with CIN with apparent transition areas. This circumstantial evidence gives biological plausibility to the suggestion that HPV may be a cervical carcinogen.
  - label: supports_H1
- PubMed ID: 6282442
  - evidence: Histologic HPV infection was found in 73/80 women (91%) with cervical neoplasia versus 10/80 matched controls (12.5%). The strength, specificity, and consistency of the relationship suggest SPI may be a precursor of cervical malignancy.
  - label: supports_H1
- PubMed ID: 6086470
  - evidence: HSV-2-specific antibodies were found in 24% of cervical-cancer patients and 17% of controls. The abstract states that a causal relationship between HSV-2 and cervical cancer has never been proven.
  - label: inconclusive
- PubMed ID: 6099637
  - evidence: Repeated intravaginal inoculation of mice with inactivated HSV-2 induced cervical carcinoma in approximately 50% of mice. Prior HSV-1 subunit-vaccine immunization reduced cervical-carcinoma frequency to 19% in this mouse model.
  - label: supports_H2
- PubMed ID: 6277023
  - evidence: The reported HSV-2 evidence is described as albeit circumstantial, and HSV-2-induced cervical atypia would have to be very rare outcome resulting from a very common exposure. The abstract proposes that papillomavirus and other sexually transmitted pathogens may induce atypia or interact with HSV-2, and calls for prospective studies.
  - label: inconclusive
- PubMed ID: 6282116
  - evidence: Among women with one/no sex partner and those with two or more partners, cervical-neoplasia risk was elevated when HSV-2 antibodies were present. The authors state that the findings may strengthen credence in HSV-2 as an etiologic factor in cervical cancer.
  - label: supports_H2
- PubMed ID: 6300393
  - evidence: Studies showed a higher incidence of HSV-2 infection in cervical-cancer patients than in matched controls, and HSV markers were reported in transformed cells. HSV has transformed cells in culture, and a mouse model for virus-induced cervical cancer is described as showing promise.
  - label: supports_H2

Overall score rationale (LLM-generated):

Eight abstracts support H1, including cancer-enriched HPV16/HPV18 DNA and possible viral integration (PMID: 6304740, 6329740), a 91% versus 12.5% neoplasia-control HPV association (PMID: 6282442), and infection-to-neoplasia spectrum evidence (PMID: 6318956, 6299907). Three abstracts support H2 through an HSV-2 mouse cervical-carcinoma model, epidemiologic HSV-2 antibody associations, and reported cellular/animal oncogenic findings (PMID: 6099637, 6282116, 6300393). HSV support is tempered by abstracts stating that human causality has never been proven, that evidence is circumstantial, and that HSV-2 seroepidemiology is inconsistent; HPV is characterized as clearly more important (PMID: 6086470, 6277023, 6399280). A score of 78 is above the 75 anchor because the HPV evidence is more numerous, more consistent, and includes cancer-biopsy molecular findings, but is below 100 because the set contains substantive HSV epidemiologic and experimental support rather than direct refutation of H2.

Tallies:

- support H1: 8
- support H2: 3
- both: 0
- neither or inconclusive: 2

**Score**: 78,

Decision: H1

##### Peptic Ulcer

The following is an example of KM-GPT-DCH results for a 1987 run (iteration 1) of the cause of peptic ulcer where hypothesis 1: Peptic ulcers are caused by a bacterial infection or hypothesis 2: Peptic ulcers are caused by psychological stress. 1987 is the year that the algorithm begins to favor bacterial infection as the primary cause of peptic ulcers.

Per abstract reasoning (LLM-generated):

- PubMed ID: 3573680,
  - evidence: Campylobacter-like organisms were found within duodenal mucosa at active ulcer edges; their presence with polymorphonuclear leukocytes suggests the organisms are pathogenic.
  - label: supports_H1
- PubMed ID: 2861652
  - evidence: Reduced acid secretion permits gastric bacterial colonization; the abstract discusses bacterial overgrowth as a consequence of hypochlorhydria, not as a cause of peptic ulcer.
  - label: neither
- PubMed ID: 6357964
  - evidence: In a randomized duodenal-ulcer trial, bismuth and cimetidine produced similar healing, while cimetidine increased gastric pH and bacterial isolates.
  - label: neither
- PubMed ID: 3576129
  - evidence: C. pyloridis was cultured in all 10 patients with active gastritis and active duodenitis and was closely related to active gastritis, fulfilling a criterion for localized bacterial infection.
  - label: supports_H1
- PubMed ID: 6145023
  - evidence: Curved bacilli were present in almost all patients with active chronic gastritis, duodenal ulcer, or gastric ulcer and may be an important factor in the aetiology of these diseases.
  - label: supports_H1
- PubMed ID: 3813480
  - evidence: This is a case of abdominal actinomycosis three years after surgery for duodenal peptic ulcer; it does not address the cause of the ulcer.
  - label: neither
- PubMed ID: 4015029
  - evidence: After Billroth I surgery, gastric-remnant bacterial colonization and atrophy were described, but postoperative gastritis did not correlate with duodenogastric reflux.
  - label: neither
- PubMed ID: 3623220
  - evidence: Ranitidine and vagotomy increased gastric pH, nitrate, and bacterial counts, but an acid tide prevented persistent gastric colonisation during ranitidine treatment.
  - label: neither
- PubMed ID: 3582869
  - evidence: C. pyloridis was detected in 19/21 duodenal-ulcer patients, and its association with duodenal ulcers was statistically significant (p<0.001).
  - label: supports_H1
- PubMed ID: 4084017
  - evidence: Cimetidine moderately inhibited neutrophil bactericidal activity in duodenal-ulcer patients; the study does not assess bacterial or psychological causation of ulcers.
  - label: neither
- PubMed ID: 3587292
  - evidence: No abstract available.
  - label: inconclusive
- PubMed ID: 3297911
  - evidence: GCLOs were associated with gastritis and peptic ulceration, but the abstract states their pathogenetic significance was unknown and whether they cause, colonize, or worsen gastritis was unanswered.
  - label: inconclusive
- PubMed ID: 4073078
  - evidence: This randomized study concerns antibiotic treatment of skin and soft-tissue infections; the single bleeding peptic-ulcer death does not address ulcer causation.
  - label: neither
- PubMed ID: 3982344
  - evidence: No abstract available.
  - label: inconclusive
- PubMed ID: 3652784
  - evidence: C. pyloridis culture was positive in 77% of gastric-ulcer and 84% of duodenal-ulcer cases, compared with 53% of active chronic gastritis cases.
  - label: supports_H1
- PubMed ID: 3535017
  - evidence: Cimetidine increased total and nitrate-reducing gastric bacteria, whereas colloidal bismuth did not; the report concerns hazards of acid suppression rather than ulcer causation.
  - label: neither
- PubMed ID: 4032128
  - evidence: Mucosa-related bacteria occurred in about 80% of peptic-ulcer patients and rarely in controls; bacterial numbers correlated positively with intraepithelial polymorphonuclear leukocytes.
  - label: supports_H1
- PubMed ID: 3950448
  - evidence: Spiral bacteria were found in 69% of gastritis specimens versus 14% of histologically normal specimens; 10/17 gastric-ulcer patients also had spiral bacteria.
  - label: supports_H1
- PubMed ID: 3310428
  - evidence: In active peptic-ulcer disease, C. pylori was found in duodenal mucosa, including intercellular spaces and epithelial vacuoles; these findings raise a hypothesis of direct mucosal damage.
  - label: supports_H1
- PubMed ID: 6703665
  - evidence: Multiple mucosal bacterial species were isolated from peptic-ulcer patients, and ampicillin reduced their adhesion before surgery; no link to ulcer causation was established.
  - label: neither
- PubMed ID: 3099375
  - evidence: Spiral bacteria were detected in gastric specimens and described as supposed to be aetiologically related to peptic ulcer and gastritis, but the abstract provides no causal test.
  - label: inconclusive
- PubMed ID: 6343664
  - evidence: This randomized trial compares cefotetan and cefmetazole for purulent peritonitis, including perforated ulcers, not the causes of peptic ulcer disease.
  - label: neither
- PubMed ID: 3615089
  - evidence: C. pyloridis was associated with primary antral gastritis in children, but the abstract does not establish an association with or cause of childhood peptic ulcer.
  - label: neither
- PubMed ID: 6698439
  - evidence: After operations for duodenal ulcer, higher pH and bacteria were sometimes observed, but there was no increased total N-nitrosocompound exposure.
  - label: neither
- PubMed ID: 6740211
  - evidence: Pernicious-anaemia patients had bacterial overgrowth, duodenitis, and dysplasia; this does not evaluate bacterial or psychological causes of peptic ulcer.
  - label: neither
- PubMed ID: 3832145
  - evidence: Alexithymia was detectable in an unselected psychosomatic group that included duodenal-ulcer patients, with high scores in male psychosomatic patients.
  - label: supports_H2
- PubMed ID: 3770362
  - evidence: In a controlled study, ulcer patients reported more negative perceptions of life events, personality disturbances, fewer supports, depression, and anxiety than controls.
  - label: supports_H2
- PubMed ID: 3625586
  - evidence: Compared with community controls, gastric-ulcer patients had higher trait anxiety and psychoticism, while the duodenal-ulcer group had higher state anxiety.
  - label: supports_H2
- PubMed ID: 6523076
  - evidence: Chronic duodenal-ulcer patients showed high anxiety, dependence, and introversion; anxiety was higher among patients with elevated acid-output and/or pepsinogen measures.
  - label: supports_H2
- PubMed ID: 6834368
  - evidence: Patients linked indigestion chiefly with psychological factors, feeding patterns, and bowel function; this terminology study does not establish a cause of peptic ulcer.
  - label: neither
- PubMed ID: 3773295
  - evidence: Among occupational patients, the duodenal-ulcer group had greater work stress than the gastric-ulcer group; smoking and stressful work environment were correlated.
  - label: supports_H2
- PubMed ID: 3784513
  - evidence: No abstract available.
  - label: inconclusive
- PubMed ID: 3963250
  - evidence: Former prisoners of war had a higher rate of postwar duodenal ulcer than non-POW combatants, but the abstract does not link individual psychological measures to ulcer occurrence.
  - label: inconclusive
- PubMed ID: 6678135
  - evidence: In 30 duodenal-ulcer patients, recent-life-event stress was low and anxiety, depression, smoking, and drinking were not more common than in the general population.
  - label: inconclusive
- PubMed ID: 6506980
  - evidence: Functional psychosomatic gastrointestinal disorders followed psychotrauma, but comparison with duodenal ulcer noted that these personality features tended to decrease with further somatization.
  - label: inconclusive
- PubMed ID: 3895386
  - evidence: In a randomized trial, adding psychotherapy to medical treatment improved outcomes for peptic-ulcer patients at 15 months compared with medical treatment alone.
  - label: supports_H2
- PubMed ID: 3726009
  - evidence: Trait psychological indices in endoscopically identified ulcer patients may be of causal significance; duodenal-ulcer patients had higher introversion and psychoticism than controls.
  - label: supports_H2
- PubMed ID: 3091469
  - evidence: This review finds agreement that peptic-ulcer patients have dependence/independence problems and high anxiety, while evidence on stress is described as nonuniform and often contradictory.
  - label: supports_H2
- PubMed ID: 3628682
  - evidence: In a matched case-control study, chronic gastric-ulcer patients were characterized by anxiety, dependence, introspection, trustfulness, and adaptability.
  - label: supports_H2
- PubMed ID: 3615786
  - evidence: This study compares affect expression in irritable-bowel, duodenal-ulcer, and anxiety-disorder patients; it does not test whether psychological stress causes ulcer disease.
  - label: neither
- PubMed ID: 6677254
  - evidence: No abstract available.
  - label: inconclusive
- PubMed ID: 3765911
  - evidence: In recurrent duodenal ulcer, stress-related bereavement and nonspecific activation were typical of recidivation; long-term prophylaxis included psychopharmacological and psychotherapeutic approaches.
  - label: supports_H2
- PubMed ID: 3949118
  - evidence: Duodenal-ulcer subjects were more neurotic, anxious, and depressed than community controls, although the reported numerical differences were small and correlations low.
  - label: supports_H2
- PubMed ID: 4068632
  - evidence: No abstract available.
  - label: inconclusive
- PubMed ID: 3477993
  - evidence: The review states that emotional stress can increase gastric secretion, but says the role of psychosomatic reactions in organic diseases such as peptic ulcer remains unclear.
  - label: inconclusive
- PubMed ID: 6542917
  - evidence: Duodenal-ulcer patients had distinct electrodermal responses, and anxiety and depression scores were higher in both psychosomatic patient groups than in controls.
  - label: supports_H2
- PubMed ID: 4080928
  - evidence: This childhood review states that certain psychological stresses may play a significant role in peptic-ulcer pathogenesis in genetically predisposed children.
  - label: supports_H2
- PubMed ID: 3895389
  - evidence: Among 103 peptic-ulcer patients, authors considered psychological factors often significant and proposed IBS and peptic ulcer as facets of an underlying psychogenic mechanism, while noting conclusive evidence was lacking.
  - label: supports_H2

Overall score rationale (LLM-generated):

Eight abstracts provide disease-specific bacterial evidence: bacteria were strongly associated with duodenal or gastric ulcers (PMID: 3582869, 3652784, 4032128, 3950448) and were observed invading or damaging ulcer-associated mucosa (PMID: 3573680, 3310428). The bacterial case is limited because much of the evidence is observational association, and one review explicitly states that GCLO pathogenetic significance and whether they cause disease remained unknown (PMID: 3297911).

Fourteen abstracts support psychological involvement, including controlled associations with stress, personality traits, anxiety, or depression (PMID: 3770362, 3625586, 3726009, 3628682, 3949118) and improved peptic-ulcer outcomes when psychotherapy was added to medical therapy in a randomized trial (PMID: 3895386). Psychological evidence is also substantially associative and mixed: stress evidence was described as contradictory (PMID: 3091469), organic-disease causation as unclear (PMID: 3477993), and one duodenal-ulcer pilot found low recent-life-event stress (PMID: 6678135).

Applying the continuous anchors, the direct, disease-localized bacterial findings modestly outweigh the larger but less causally specific and mixed psychological literature; thus the score is moderately above 50 rather than near the strong-support anchor of 100.

Tallies:

- Support H1: 8
- Support H2: 14
- Both: 0
- Neither or inconclusive: 26

score: 61,

decision: H1

##### ASCVD

Finally, we show an example KM-GPT-DCH run for the competing hypotheses for preventing atherosclerotic cardiovascular disease (ASCVD), statins (hypothesis 1) or hormone replacement therapy (HRT) (hypothesis 2). While early data raised the hypothesis of HRT for ASCVD reduction, in 2002, use of hormone therapy to reduce ASCVD risk was refuted by randomized control trials, highlighting the rejection of this clinical hypothesis. Our algorithm demonstrates this shifting temporal trends in therapies. Early on KM-GPT-DCH supports HRT, with robust demarcation when the hypothesis was rejected, and the rising and strong support of statins starting in the 1990s. Initial longitudinal cohort observational data in 1991 [39], demonstrated a reduction in ASCVD risk (relative risk reduction 0.50 with a 95% CI, 0.43-0.56) with the use of HRT, which led to widespread adoption of use to treat menopausal symptoms with the potential to reduce ASCVD events [40]. However, this idea was subsequently refuted in two large randomized clinical trials: the Heart and Estrogen/progestin Replacement Study (HERS) in 1998 and in the Women’s Health Initiative (WHI) in 2002-2004 [41–44], leading to significant concerns about HRT use and risk of harms in post-menopausal women resulting in drastic reductions (70-80%) in the use of HRT to treat menopausal symptoms [45]. Currently, all major societies recommend against using HRT for ASCVD prevention at any age, this was further supported by a 2025 Cochrane review showing no evidence that HRT has any role in the treatment or prevention of ASCVD [46, 47]. Despite these findings, the clinical need to manage menopausal symptoms persisted. Subsequent secondary analyses suggested that the cardiovascular harms of HRT may be time-dependent, with lower risks when initiated closer to menopause onset, informing current guidelines that support risk/benefit discussions for HRT use for menopausal symptoms in eligible, recently menopausal individuals [45, 48, 49]. Currently, in clinical practice, HRT is used primarily to address moderate-severe vasomotor and genitourinary symptoms of menopause in eligible individuals, with therapy individualized based on discussion of risks and benefits [50].

Example KM-GPT-DCH result from 2002 (iteration 3):

Per abstract reasoning (LLM-generated):

- PubMed ID: 11928058
  - evidence: EuroASPIRE II used statins more frequently, but secondary prevention has not substantially improved\ between 1996 and 2000.
  - label: neither
- PubMed ID: 11901897
  - evidence: The review notes reduced C-reactive protein with a statin, but does not report cardiovascular-prevention outcomes for statins.
  - label: neither
- PubMed ID: 10956605
  - evidence: Guidelines state there is important evidence for successful lipid control with lipid-lowering drugs, especially statins, and identify statin lipid control among the most efficient measures to reduce morbidity and mortality.
  - label: supports_H1
- PubMed ID: 11501602
  - evidence: Despite the efficacy of statin therapy in reducing the incidence of CHD, treated patients may still experience CHD events.
  - label: supports_H1
- PubMed ID: 11769362
  - evidence: The Stockholm survey reports marked undertreatment and projects costs if statin recommendations are implemented, without reporting cardiovascular outcomes.
  - label: inconclusive
- PubMed ID: 12197410
  - evidence: This pathogenesis review says statins lower CRP and may lower atherosclerotic risk beyond cholesterol lowering, but provides no cardiovascular outcome comparison.
  - label: inconclusive
- PubMed ID: 11282901
  - evidence: In a randomized placebo-controlled trial, fluvastatin reduced common-carotid IMT progression at 36 months (-0.009 mm/y; P=0.002).
  - label: supports_H1
- PubMed ID: 11347831
  - evidence: The study concerns statins and steroid-associated osteonecrosis; it does not evaluate cardiovascular-disease prevention.
  - label: neither
- PubMed ID: 11501230
  - evidence: The abstract states that effective hyperlipidaemia treatment can prevent CHD development and reduce cardiovascular events and mortality; it presents rosuvastatin as a potent statin.
  - label: supports_H1
- PubMed ID: 9694549
  - evidence: In vitro and animal-model findings suggest direct arterial-wall antiatherosclerotic effects of statins that \could translate\ into cardiovascular prevention.
  - label: inconclusive
- PubMed ID: 11688065
  - evidence: Combined treatment of hyperglycaemia, dyslipidaemia, and hypertension is said to decrease diabetic complications, but the study does not isolate a statin effect on cardiovascular prevention.
  - label: inconclusive
- PubMed ID: 9597427
  - evidence: Long-term HMG-CoA reductase inhibitor treatment is described as effective, safe, and tolerable with 30% mortality reduction; it is recommended as cost-effective for secondary prevention.
  - label: supports_H1
- PubMed ID: 10980827
  - evidence: Preliminary data suggest that the relative efficacy of statins and aspirin in secondary prevention may depend on baseline CRP, without reporting a definitive statin outcome effect.
  - label: inconclusive
- PubMed ID: 11151651
  - evidence: The Health Council recommends lipid-lowering therapy, particularly statins, for atherosclerotic disease and other markedly high-risk groups.
  - label: supports_H1
- PubMed ID: 11452674
  - evidence: No abstract available.
  - label: neither
- PubMed ID: 11829452
  - evidence: Large diabetic lipid-lowering trials of statins were ongoing; the abstract says their results may determine future management.
  - label: inconclusive
- PubMed ID: 12006393
  - evidence: Despite major lipid reductions with atorvastatin, intensive statin therapy showed no effect on NO-dependent or endothelium-independent forearm vasodilation in type 2 diabetes.
  - label: neither
- PubMed ID: 12394655
  - evidence: In hypercholesterolemic people, statin therapy plus Mediterranean diet was associated with lower acute coronary syndrome risk (OR 0.57; P<0.01), though the authors state causal evidence cannot be claimed.
  - label: supports_H1
- PubMed ID: 11240415
  - evidence: Statins are recommended for diabetes with hypercholesterolaemia, but their efficacy for typical diabetic dyslipidaemia and cardiovascular prevention \remains to be demonstrated.
  - label: inconclusive
- PubMed ID: 11798151
  - evidence: For dialysis patients, real benefit from lipid-lowering has not yet been demonstrated and randomized statin trials were underway.
  - label: inconclusive
- PubMed ID: 12006916
  - evidence: Landmark statin trials are said to demonstrate unequivocally that effective lipid-lowering therapy significantly decreases CHD morbidity and mortality.
  - label: supports_H1
- PubMed ID: 10944975
  - evidence: For cardiovascular prevention, statins are identified as first-line drugs for high cholesterol; secondary prevention includes LDL targets and drug treatment thresholds.
  - label: supports_H1
- PubMed ID: 11978147
  - evidence: Among MI survivors, statin use was associated with lower recurrent coronary events (RR 0.59) and any atherosclerotic cardiovascular event (RR 0.63) versus no lipid-lowering treatment.
  - label: supports_H1
- PubMed ID: 12505222
  - evidence: Children with familial hypercholesterolaemia randomized to simvastatin had significantly improved flow-mediated dilation versus placebo, toward control levels.
  - label: supports_H1
- PubMed ID: 11245405
  - evidence: Clinical trials have demonstrated that fatal and nonfatal coronary events and strokes can be prevented; statins are described as commonly used to lower cholesterol in cardiovascular prevention.
  - label: supports_H1
- PubMed ID: 12632840
  - evidence: In the randomized PROSPER trial, pravastatin lowered LDL 34% and reduced the composite coronary death, MI, or stroke endpoint by 15% (P=0.014).
  - label: supports_H1
- PubMed ID: 12467937
  - evidence: For combined dyslipidaemia, statin monotherapy has been shown to benefit these patients, although optimal combination therapy still required trials.
  - label: supports_H1
- PubMed ID: 11282915
  - evidence: The review states that aspirin and statins seem to attenuate inflammatory risk and that hsCRP may help target proven primary-prevention therapies.
  - label: supports_H1
- PubMed ID: 9529266
  - evidence: In a 3-year placebo-controlled trial, postmenopausal hormone therapy produced sustained 17%-23% reductions in lipoprotein(a), a cardiovascular risk factor.
  - label: supports_H2
- PubMed ID: 10448608
  - evidence: Although observational studies suggested benefit, the randomized HERS trial failed to demonstrate cardioprotection and showed early adverse outcomes in women with coronary disease.
  - label: neither
- PubMed ID: 10764660
  - evidence: Clinically used estrogens inhibited cultured human aortic smooth-muscle-cell growth, migration, collagen synthesis, and MAP kinase activity through estrogen receptors.
  - label: inconclusive
- PubMed ID: 10746517
  - evidence: Women using HRT had improved lipid profiles and iron-status parameters; the authors say these findings suggest a need to further examine the relation to decreased cardiovascular risk.
  - label: inconclusive
- PubMed ID: 10795496
  - evidence: The randomized HERS study showed increased cardiovascular events in year 1 and a later decrease, with \no overall effect.
  - label: neither
- PubMed ID: 11282054
  - evidence: HERS failed to demonstrate benefit, and the review concludes that an evidence-based clinical decision on estradiol for cardiovascular prevention was not then possible.
  - label: inconclusive
- PubMed ID: 11795349
  - evidence: Raloxifene and HRT favorably alter some cardiovascular risk factors, but proof that either reduces heart-disease risk must await cardiovascular endpoint trials.
  - label: inconclusive
- PubMed ID: 12186518
  - evidence: An adjusted meta-analysis found no CAD-incidence benefit with HRT (RR 1.07) and was consistent with randomized trials showing no primary or secondary prevention benefit.
  - label: neither
- PubMed ID: 10549564
  - evidence: This review lists beneficial cardiovascular effects among current health benefits associated with hormone replacement therapy.
  - label: supports_H2
- PubMed ID: 10717485
  - evidence: The review states that studies documented menopause-associated cardiovascular risk and risk reduction in this area by HRT.
  - label: supports_H2
- PubMed ID: 10348700
  - evidence: HRT significantly decreased lipoprotein(a) and PAI-1 and increased fibrinolytic activity in postmenopausal women, mechanisms relevant to cardiovascular risk.
  - label: supports_H2
- PubMed ID: 11821592
  - evidence: These HRT guidelines conclude that therapy protects against osteoporosis and \reduces the risk of cardiovascular disease.
  - label: supports_H2
- PubMed ID: 12435256
  - evidence: In the randomized WAVE trial in women with coronary disease, HRT provided no cardiovascular benefit; inclusion of death or MI showed increased risk with active HRT (P=0.045).
  - label: neither
- PubMed ID: 11004936
  - evidence: The abstract only says the article discusses clinical data, recommendations, and future directions for HRT and cardiovascular prevention; no results are provided.
  - label: neither
- PubMed ID: 11466136
  - evidence: HRT shortened QT dispersion in healthy postmenopausal women, but the abstract says the clinical implications must be clarified.
  - label: inconclusive
- PubMed ID: 10566666
  - evidence: HRT substantially improved microvascular reactivity in healthy postmenopausal women, whereas benefit was less apparent in women with type 2 diabetes.
  - label: supports_H2
- PubMed ID: 11431642
  - evidence: The review asserts that postmenopausal HRT prevents cardiovascular disease, while also noting uncertainty surrounding evidence and risks.
  - label: supports_H2
- PubMed ID: 11521121
  - evidence: The review describes protection against cardiovascular disease as an emerging HRT benefit, while noting a need to balance tolerability and risks.
  - label: supports_H2
- PubMed ID: 11139791
  - evidence: HERS failed to reduce subsequent events in postmenopausal women with CAD and instead reported an early increase in CAD events; the authors advise against starting HRT solely for CAD prevention after a recent event.
  - label: neither
- PubMed ID: 9842237
  - evidence: The review discusses hormone replacement therapy benefits in prevention of cardiovascular disease and postmenopausal osteoporosis.
  - label: supports_H2
- PubMed ID: 9798134
  - evidence: The article says HRT is likely to become important for protection from increasing cardiovascular risk, but the reported placebo-controlled patch trials evaluated climacteric symptoms rather than cardiovascular events.
  - label: inconclusive
- PubMed ID: 10900674
  - evidence: Despite reported anti-ischemic mechanisms and functional findings, HRT for cardiovascular reasons in coronary disease is considered ambiguous and cannot generally be recommended for secondary prevention.
  - label: inconclusive

Tallies:

- support_H1: 17
- support_H2: 6
- both: 1
- neither or inconclusive: 3

Overall score rationale (LLM-generated):

Robust RCTs and multiple reviews (PubMed IDs 12114036, 12523144, 12817199) consistently confirm statins lower MI, stroke, and mortality. 17 abstracts support H1, only 6 support H2. One citation (15765625) explicitly reports increased CV risk with HRT, tempering Hormone-therapy evidence; others are observational or mechanistic. Overall evidence is roughly triple in volume and higher quality for statins versus hormone therapy, aligning with guideline recommendations (12448270).

score: 78

decision: H1

#### Additional information on controversial hypotheses

##### Peptic ulcer caused by bacteria hypothesis

The first description of a spirochete and suggestion of its role in gastric ulcers was first mentioned in 1875 by Bottcher and Letulle [51]. Warren first noted the spirochetes associated with gastric ulcer in 1979, isolated them in 1982, and published the results in 1983 [52]. *Helicobacter pylori* was originally called *Campylobacter pyloridis* in 1982, changed to *Campylobacter pylori* in 1987 [53], and finally reclassified as *Helicobacter pylori* in 1989 [54].

Approximately 10% of peptic ulcer cases are idiopathic. It is possible that stress or some other genetic or infectious cause could be primary in these cases.

##### Scrapie caused by prions hypothesis

The causal agent of scrapie has long been debated. An early proposal was the slow virus hypothesis [55], followed by the virino hypothesis proposed in the late 1970s and early 1980s [56] followed by prions being a distinct infectious agent [57]. Note that the self-replicating protein idea was first proposed by Griffith, with support from experiments by Alper and Pattinson in 1967 [58]. Support for the prion hypothesis strengthened during the 1990s [59]. In recent years, many other neurodegenerative diseases are being termed prion-like (such as Alzheimer's disease, ALS) having the alteration in protein conformation as an important contributor or underlying cause [60, 61].

##### Genetic predisposition, vaccines, and autism

While scientific evidence supports the genetic predisposition theory but does not support the vaccine theory of autism, controversy persists in the public. Despite the evidence against the vaccine hypothesis and for the genetic predisposition hypothesis, childhood vaccination rates have been decreasing in the US [62]. Indeed, numerous factors likely contribute to portions of the public continuing to endorse the vaccine-autism theory despite the existing scientific evidence for genetic predisposition. For one, the gene theory of autism is complex, suggesting that the genetic susceptibility of autism stems from a combination of multiple sources (*e.g.*, common inherited variation, rare inherited genes, and rare *de novo* genes) [63–66] that may contribute differently to autistic traits, motor skills, and intellectual disability [63, 67, 68]. Because of the gene-theory complexities, people may prefer the simpler, albeit less data-based theory, of vaccines and autism. Additionally, the movement to democratize health decisions by eschewing expertise may contribute to people rejecting the scientific consensus [69]. However, there remains no link between autism and vaccines (for a systematic review and meta-analysis see [70]). Meanwhile, the evidence for genetic contributions and their ability to account for the considerable heterogeneity in the autism behavioral profile [71] and neurobiological profile [72] continues to grow.

#### Current competing hypotheses

Additional detail on the 20 currently controversial and/or competing hypotheses is provided here, as well as in **Supplementary Table 4**.

##### Alzheimer’s disease

Alzheimer's disease (AD) is characterized by severe dementia and memory loss, and is thought to be caused by either the accumulation of a mutated form of the amyloid‑beta (Aβ) protein in the brain that results in a plaque (H1) or the hyperphosphorylation of the tau (τ) protein that then aggregates and forms neurofibrillary tangles (H2) [73]. In support of the Aβ plaque hypothesis, mouse models show a reduction of Aβ plaques by various drugs improved cognitive abilities [74, 75]. Additionally, in autosomal dominant Alzheimer's disease (ADAD), Aβ plaques accumulate higher in ADAD carriers of the gene than non-carriers, and the plaques accumulate before cognitive impairment occurs [76]. In the tau protein hypothesis, this microtubule binding protein undergoes hyperphosphorylation which causes the dissociation of tau from microtubules, leading to neurofibrillary tangles and the onset of AD [77]. It has been shown that tau-targeted immunotherapies can rescue cognitive deficits in multiple animal or clinical studies [78–80]. Additionally, a longitudinal human study shows that tau accumulation is associated with memory impairment independent of Aβ status, indicating tau proteins may play a more primary role in AD [81]. Finally there is some evidence of a synergistic interaction between Aβ and tau, implicating both hypotheses as causal [82, 83]. Our model finds about twice as many abstracts supporting tau pathology as being the primary cause, compared to abstracts that support the amyloid plaque hypothesis. Overall, our model gives a posterior mode score of 37.6 with the credible interval just shy of the 50 mark, so the tau pathology hypothesis is slightly more favored.

##### Endometriosis

What is the mechanism of the pathogenesis of endometriosis? Endometriosis develops when endometrial tissue grows outside of the uterus, causing lesions, pain, nodules, infertility, and/or other symptoms [84, 85]. Two of the main competing theories for the pathogenesis of this are retrograde menstruation (H1) and coelomic metaplasia (H2). Retrograde menstruation happens when menstrual blood containing endometrial cells flows backward into the peritoneal cavity and then these cells then implant and grow lesions [85]. However, this method does not explain the rarer occurrences of endometriosis that happens outside of the peritoneal cavity. The coelomic metaplasia theory, on the other hand, suggests that coelomic epithelial cells transdifferentiate (metaplasia) and form endometrial stroma. This would explain endometriosis that occurs in other places, such as the ovary [85]. Our model leans toward the main causal hypothesis being retrograde menstruation (with a posterior mode score of 83.3), and most of the abstracts support this hypothesis, but it is possible different types of endometriosis exist and have different causes.

##### REM sleep

What is the primary function of REM sleep? REM sleep is characterized by rapid eye movement, enhanced brain activity, and often dreaming, and oscillates between non-REM sleep 4-5 times per night [86]. The purpose of REM sleep is debated, with two main hypotheses being memory consolidation (H1) and brain homeostasis (H2). Memory consolidation involves the coding and storage of important memories [86]. The evidence for this occurring during REM sleep includes inability to recall recent memories that have not already been learned after REM sleep deprivation [87]. The counter-argument of brain homeostasis is that REM sleep is essential for emotional well-being and optimizing stress response, and REM sleep abnormalities are linked to psychiatric disorders like depression and post-traumatic stress disorder [87, 88]. Our model leans heavily toward the hypothesis of memory consolidation (posterior mode score 83.6) with most literature supporting this hypothesis; however, it is possible that both functions could be happening together.

##### Red wine

Cardiovascular health benefits have been associated with red wine [89], but it is unclear as to whether this is due to antioxidants present in the skin of grapes or due to the alcohol itself [90]. Polyphenol antioxidants in red wine, such as resveratrol, have been shown to have therapeutic benefits to conditions such as pulmonary arterial hypertension and reduce oxidative stress [89, 91], and alcohol abuse can have negative cardiovascular effects [92]. However, alcohol in moderation is thought to have an antithrombotic effect through the inhibition of platelet function and also decrease in LDL cholesterol concentration, both of which help prevent cardiovascular disease [93]. With a posterior mode score of 78.3, our algorithm finds most papers support the antioxidant hypothesis.

##### Polycystic Ovary Syndrome

What is the driver of polycystic ovary syndrome (PCOS)? PCOS is a common infertility disorder and symptoms include hyperandrogenism and polycystic ovaries [94]. It is not well known what the underlying cause is for PCOS, whether it is primarily an ovarian or a neuroendocrine abnormality. The ovarian hypothesis states that defects begin in the ovary and affect androgen production while the neuroendocrine hypothesis indicates that abnormalities in the brain cause the ovary dysfunction. In support of the ovarian hypothesis, when compared to normal theca cells (TCs) from the ovaries, TCs from PCOS patients produce androgens at a higher rate, causing hyperandrogenism, which in turn causes a decrease in autophagy, which can ultimately cause PCOS [95]. In support of a neuroendocrine cause, women with PCOS have been shown to have frequent [gonadotropin-releasing hormone](https://en.wikipedia.org/wiki/Gonadotropin-releasing_hormone) (GnRH) pulses, which is a neurohormone from the pituitary gland in the brain, and higher levels of the signaling molecule kisspeptin, which initiates GnRH secretion [96]. Our algorithm has mixed support for both causes with a posterior mode score of 53.5, and relevant abstracts cite both central neuroendocrine dysfunction and ovarian dysfunction.

##### Aging

What is the primary mechanism of aging? The first hypothesis is that aging is the result of stochastic damage to the DNA and cells that accumulates over time. The second hypothesis is that specific genes regulate changes and hormonal signaling to induce aging, likely to limit population size and induce generational turnover. In support of the first hypothesis that DNA damage causes aging, it has been shown that deletions in mitochondrial DNA correlate with aging as well as other disorders [97]. Additionally somatic mutations throughout the body accumulate with age [98, 99] and DNA damage induces senescence in cells [100]. However, loss of function of specific gene products (like parathymosin) have been shown to cause neurodegeneration and a reduced lifespan [101] and genome-wide association studies have linked multiple loci with longevity [102], supporting the second genetic hypothesis. Our algorithm favors the stochastic DNA damage hypothesis as the primary cause of aging with a posterior mode score of 70.7 because this has been shown in a mechanistic way, and more publications support this hypothesis.

##### Acupuncture

How does acupuncture treatment produce analgesic effects? The first hypothesis attributes the analgesic effects to the release of endorphins when needling [103]. The second hypothesis argues that pain is alleviated due to diffuse noxious inhibitory controls (DNIC), in which a counter irritation inhibits wide dynamic range (WDR) neurons, which in turn inhibits the pain [104]. Studies have found that acupuncture increases β-endorphin levels [105, 106], supporting hypothesis one. Some evidence for the DNIC mechanism (hypothesis two) exists as well in rats where electro-acupuncture suppressed the firing of C-fiber-evoked WDR neurons [107]. However, the number of studies that link pain-relief from acupuncture to increases in endorphins include clinical trials and outnumber studies with supporting DNIC as the causal mechanism, and our algorithm favors the first hypothesis with a posterior mode score of 84.9.

##### Cancer Progression

What is the primary reason for the progression of cancer? Two main competing hypotheses in the field include the Somatic Mutation Theory (SMT) and the Tissue Organization Field Theory (TOFT) [108]. The SMT argues that cancer begins with a single cell that accumulates mutations in its DNA which allow it to replicate uncontrollably, and this initiates and drives cancer progression [109]. Evidence for SMT includes the sequencing of several different types of tumors and finding hundreds of cancer driver genes with mutations under positive selection that cause the progression of cancer [110]. Additionally many different types of cancer identify somatic mutations in tumor suppressor or oncogenes that facilitate the progression of cancer, supporting SMT [111–115]. In contrast, the tissue organization theory (TOFT) posits that the default state of cells is proliferation with variation and motility, but multicellular differentiated somatic cells are constrained by cell-cell interactions within a tissue and ultimately cannot contribute to the organism’s reproduction. When these constraints are disrupted, they return to the default state of proliferation, undergo uncontrolled replication, and cause cancer [108] [116]. Additionally, tissue organization can become dysregulated by changes in bioelectricity which exists from the cell membrane potential. A study that supports this shows that cell depolarization can cause a change in cell state and can propagate this state to other cells without any genomic mutation [117], providing evidence for TOFT. Our model reasons that the SMT theory is more likely because SMT-supporting papers span multiple tumor types and methodologies (genomics, evolutionary analyses) and have a higher number of papers supporting it, whereas TOFT support is largely conceptual or from mechanical studies, ultimately giving a posterior mode score of 77.7.

##### Cancer Origin

What is the primary origin of cancer? Whether cancer originates from cancer stem cells or through stochastic clonal evolution is currently being debated. Cancer stem cells are tumor cells that have stem-cell like properties such as the ability to self-renew, resistance to cytotoxicity, and multipotency which are thought to initiate cancer [118]. Cancer stem cells have been identified for many different types of cancers (breast cancer, lung cancer, gastric cancer, ovarian cancer, etc.) characterized by specific surface markers, enhanced DNA repair capacity, and plasticity that both initiate these cancers and help the tumors evade treatment [119–123], supporting the cancer stem cell theory. In contrast, stochastic clonal evolution is similar to the somatic mutation theory, but in addition once a mutation occurs in a somatic cell, it may incur a selective advantage on the cell resulting in proliferation and cancer, therefore the origin of cancer occurs through positive selection of these cells [124]. Evidence for this theory includes instances of tumor heterogeneity that have arisen through distinct clonal populations, consistent with positive selection of individual cells [125, 126]. Additionally, selective resistance of clonal populations to apoptosis supports that stochastic mutations driving this resistance are under positive selection and allow for the progression of cancer [127]. With a posterior mode of 56.8, our model has slightly higher support for the cancer stem cell theory, but because the credible interval crosses the 50 mark, we deem support for both theories equally valid.

##### Pre-eclampsia

What causes the pathogenesis of pre-eclampsia? Two leading theories are that the pathogenesis originates from the placenta or the maternal cardiovascular system. Pre-eclampsia (PE) is characterized by hypertension and significant proteinuria on or after 20 weeks of pregnancy, and is a leading contributor to maternal mortality [128]. The placental hypothesis indicates that there is dysfunction in placental homeostasis, where there is inadequate trophoblast invasion (connection between the placenta and uterus) and thus incomplete remodeling of uterine spiral arteries. This leads to placental ischemia, hypoxia and thus oxidative stress, and then hypertension later in the pregnancy [128, 129]. Single cell and spatial transcriptomics and metabolomics has revealed that PE placentas show an increase in placental macrophages, enrichment of transcriptomic pathways related to oxidative stress, and dysregulation of lipid and energy metabolism, supporting a placenta dysfunction hypothesis [129]. Additionally, other studies show gene or protein levels linked to trophoblast invasion from the placenta are altered in PE placentas, supporting the hypothesis that the placental defects are the main cause of PE [130, 131]. In contrast the maternal cardiovascular hypothesis posits that there is a concurrent maladaptation of the maternal cardiovascular system along with abnormal placentation, which predisposes the mother to PE [132]. A large study (including > 187,000 women) found that women with pregnancy complications such as PE had many cardiometabolic risk factors, such as higher blood pressure, glucose levels, and body mass indices, 10 years before their pregnancy compared to women without complications [133]. An additional study linked high cardiovascular health with a lower risk of PE [134]. This supports that poor cardiometabolic health before pregnancy (and thus the development of the placenta) may contribute to developing PE. Some papers suggest that early and late onset of preeclampsia may have different causes, the early onset being the placenta-dominant form while the late onset being due to maternal cardiac maladaptation [135]. Our model finds more papers associated with placental defects as a cause of PE than cardiovascular dysfunction but acknowledges that cardiovascular health can be a contributing factor, giving a posterior mode score of 81.0, which is in favor of the placental hypothesis.

##### Multiple Sclerosis

How does Multiple Sclerosis (MS) develop? MS is a chronic inflammatory disease that is characterized by demyelination of the central nervous system (CNS) [136]. Two main hypotheses exist for the cause and development of MS, one is that it is an autoimmune disorder, where the body’s own immune cells attack healthy myelin (the protective sheath around nerve fibers), while the other is that it is caused by the Epstein-Barr virus, which then triggers the immune system [136]. A study finding elevated levels of specific immune-related proteins had causal effects on risk of MS and that may be targeted to treat MS supports the hypothesis that it is an autoimmune disease [137]. Other evidence includes elevated levels of pro-inflammatory cytokines in response to dysregulated T-cells attacking the CNS in MS patients, and how treating inflammation through exercise can help MS symptoms [138]. In contrast, other studies claim that MS is initiated by a viral attack by the Epstein-Barr virus (EBV), where antigens from EBV mimic host antigens and cause dysregulation of T-cells and B-cells, which then results in autoimmunity [139]. Our model finds more references that support an autoimmunity causality that also detail immune-mediated mechanisms, while there are fewer abstracts supporting EBV as causal, and these are more based on associations. The model strongly tilts toward autoimmunity as the main cause of MS with a posterior mode score of 71.7.

##### Type 2 Diabetes

What is the primary defect in Type 2 Diabetes? Type 2 Diabetes mellitus (T2DM) is a chronic disease with high rates of morbidity that is characterized by a progressive loss of insulin secretion by beta cells in the pancreas [140, 141]. It is unclear as to whether T2DM is ultimately caused by peripheral insulin resistance in tissues or pancreatic beta-cell failure. In the pancreatic beta cell failure hypothesis, due to high levels of glucose in the blood, pancreatic beta cells attempt to increase insulin secretion to maintain normal levels, but cannot secrete enough insulin, and their function declines. This leads to reduced glucose uptake by target tissues, hyperglycemia, and insulin resistance [141]. Therefore beta cell dysfunction or apoptosis could be the initiating factor in T2DM, supported by the fact that many people with T2DM have a reduced beta-cell mass compared to healthy individuals [142]. However, there are some indications that insulin resistance happens before pancreatic beta cell decline. Obesity has been shown to cause insulin deficiency by producing chronic low-grade inflammation, which can impair insulin signaling leading to hyperglycemia and ultimately the development of T2DM [141], supporting an insulin resistance hypothesis. Some references state that T2DM is caused by a combination of the two hypotheses, dysfunctional pancreatic beta cells that do not secrete enough insulin and tissues not able to respond to insulin [143]. Our model acknowledges that there are an approximately even number of papers that support either hypothesis, and some that support both, so with a posterior mode score of 45.2 and a credible interval that overlaps the 50 mark, it concludes that both hypotheses are equally likely.

##### Atherosclerosis

What is the primary driver of atherosclerosis? Atherosclerosis is characterized by the build-up of plaque in the arteries, driven by the retention of apolipoprotein (apoB)-containing lipoproteins, particularly low-density lipoproteins (LDL), within the arterial wall [144]. According to the response-to-retention model, subendothelial trapping of these lipoproteins provokes a maladaptive inflammatory response that initiates and sustains plaque formation [145]. These two processes, lipid accumulation and inflammation are not competing hypotheses but interconnected mechanisms that operate synergistically [146, 147]. The causal role of LDL is supported by the fact that humans with high LDL cholesterol have an increased risk of atherosclerotic cardiovascular disease, [148] that animal studies demonstrate a high fat / high cholesterol diet causes atherosclerotic lesions[149] and that drugs like statins and other lipid lowering therapies that lower LDL cholesterol reduce the risk of cardiovascular events [150]. The role of inflammation is supported by the CANTOS trial, which demonstrated that canakinumab, an anti-inflammatory therapy targeting interleukin-1β, reduced recurrent cardiovascular events independent of lipid-level lowering [151, 152]. Additionally, patients with rheumatoid arthritis, an autoimmune disease characterized by systemic inflammation, have a 1.5- to 2-fold increased risk of coronary artery disease [153]. Our model finds that more papers between the years of 2020-2026 attribute the primary driver of atherosclerosis to inflammation than cholesterol. Our model gives a posterior mode score of 42.9, leaning toward the underlying cause of atherosclerosis to be inflammation.

This finding likely reflects temporal trends in scientific discovery rather than a true hierarchy of causation. The lipid hypothesis of atherosclerosis predates the inflammation hypothesis by decades, atherosclerosis was viewed primarily as a lipid storage disease for much of the 20th century, with the role of inflammation only gaining prominence from the 1980s onward [154]. As a result, the foundational evidence supporting the causal role of LDL in atherosclerosis is well-established and has largely been translated into clinical practice through aggressive lipid-lowering therapies [155]. Importantly, anti-inflammatory therapies for ASCVD are currently tested as adjuncts to, not replacements for, background lipid-lowering therapy [155]. The model's greater recent focus on inflammation therefore fits a landscape in which the LDL pathway is well characterized and already treated while inflammation is still being explored, and because the score of 42.9 has a credible interval that nearly overlaps 50, the model does not decisively favor inflammation and the lipid accumulation hypothesis remains supported.

##### Parkinson’s Disease

Where does Parkinson’s Disease originate? Parkinson’s disease (PD) is a neurodegenerative disease that affects the ability to move and is characterized by tremors and rigidity [156], as well as the accumulation of α-synuclein proteins in the central nervous system [157]. Competing hypotheses suggest the disease originates in the central nervous system (CNS) and spreads outward, or that the pathological proteins form in the enteric nervous system (ENS) and travel up to the brain [158]. In the CNS hypothesis, α-synuclein proteins accumulate in the amygdala or connected structure, then spread to the brain stem, and this is when motor symptoms appear [158]. Some neurons were found to be susceptible to accumulation of α-synuclein proteins because of dysfunction in their mitochondria, and mitochondrial dysfunction in the brain as well as mitochondrial gene mutations have been found in Parkison’s patients, supporting the CNS hypothesis [159]. The ENS hypothesis suggests that PD begins in the gut, and that gut dysfunction causes the marker protein α-synuclein to misfold, and then this protein travels up the vagus nerve to the brain. This is supported by the findings that misfolded α-synuclein proteins can be found in the ENS sometimes years before motor symptoms appear in PD patients, and that gut symptoms also appear before motor symptoms [156, 160]. Additionally inoculation of the α-synuclein protein to the gut wall in aged mice results in the progression of this protein to the midbrain and resulting motor symptoms [161]. Our model finds more abstracts supporting the ENS hypothesis, while there is some evidence for the CNS hypothesis, it is also proposed that there could be two separate subtypes [162]. Thus, there is more support for the ENS hypothesis, meriting a posterior mode score of 28.6.

##### Consciousness

Understanding the brain’s cognitive architecture may help unravel long-standing questions about how biological mechanisms give rise to intelligence, emotion, and behavioral decision-making. This area has gained increasing attention, as such insights could have far-reaching implications for the design of future AI systems. The Global Neuronal Workspace Theory (GNWT) (H1) proposes that the brain consists of many specialized, modular, unconscious processes, and that sufficiently strong stimuli trigger a rapid and sustained increase in neural activity that propagates from local sensory regions to a distributed network, linking frontal and posterior areas [163]. In support of the GNW theory, general anesthesia was found to decrease information integration across the brain with a sharp drop in connectivity. This suggests a breakdown in network communication, preventing the brain from broadcasting information and resulting in a transition to an unconscious state [164]. Multiple other studies [165–167] also support this theory. In contrast, the more recent Integrated Information Theory (IIT) (H2) suggests that consciousness arises from the progressive integration of domain-specific macro-consciousness emerging from numerous micro-consciousness units. In a fMRI study, it was observed that the quantification and characterization of consciousness through a measure known as Φ decreased during anesthesia as predicted by IIT [168]. Additional studies [169, 170] provide supporting evidence for IIT, and while our analysis leans towards more support for the GNW theory of neural consciousness with a posterior mode score of 59.8, the credible interval score overlaps with 50, indicating similar amounts of support for both theories.

##### Neural Code

How does the neuronal respond to stimuli and signaling responses among the network of neurons? While action potentials are primarily observed as a primary carrier of neural coding, the exact encoding mechanism in the action potential sequences is not yet resolved. This first hypothesis (H1) is the frequency or the rate that action potentials are fired, termed rate coding. This is believed to contain most of the information about the stimulus and explains muscle contractions [171]. Rate coding is robust against noise; averaging spikes over time allows the brain to ignore random errors in timing. Working memory is maintained through persistent neural firing in the prefrontal cortex even after a stimulus is no longer present, with mean firing rates directly reflecting the strength of the memory representation [172]. This supports rate coding theory, as it is the rate of neuronal discharge, not spike timing or patterns, that encodes and sustains information over time. However, the rate coding fails to explain the encoding mechanisms in case of faster reactions during visual scene changes. The competing ‘temporal coding’ hypothesis (H2) argues that precise spike timing is a significant element in neural coding [173]. While the rate coding model assumes the high frequency fluctuations in action potentials as noise, the temporal coding model suggests that they encode information. When interneurons fire within a few milliseconds of each other, their effects on Purkinje cells either combine or cancel out, so the exact timing of spikes matters for how the brain processes information, supporting the temporal coding model [174]. Additional studies [175–177] provide evidence that precise spike timing, phase, or spike order conveys key information. Our analysis finds marginally higher support towards ‘temporal coding’ with a posterior mode score of 38.3; however, the credible interval spans the 50 mark, indicating neither hypothesis is dominant.

##### Schizophrenia

Schizophrenia is a chronic, severe mental disorder often characterized by hallucinations and a distorted perception of reality, along with other symptoms such as delusions and social withdrawal [178]. Although no single known cause has been identified, leading hypotheses suggest that dysregulation in dopamine (H1) or glutamate (H2) signaling pathways may contribute to the development of schizophrenia. Studies have found schizophrenia symptoms to be associated with abnormalities with dopamine regulation, including with dopamine receptors, such as an increase in D2/D3 receptors among drug-free patients, and a decrease in these receptors in drug-treated patients [179, 180]. On the other hand, several studies [181–183] suggest that glutamate may be a primary driver of schizophrenia. These included a meta-analysis on the metabolomics of schizophrenia patients that found dysregulated glutamate/glutamine pathways related to cognitive impairment in schizophrenia spectrum disorders [181]. Our analysis suggests that both hypotheses are approximately equal in likelihood, with the posterior mode score of 53.5 and the credible interval overlapping the 50-mark line.

##### Obesity

What is the primary cause of obesity? Obesity is defined by the accumulation of excessive fat and a body mass index ≥ 30 kg/m^2^ [184]. Current hypotheses for the cause of obesity are caloric surplus/calorie imbalance, where calories in exceed calories out, or that obesity is caused by the quality of carbohydrates that are eaten, and high-glycemic carbohydrates trigger insulin that then causes fat storage. In support of the calorie imbalance hypothesis, studies in animals show that an increase in weight gain is caused by feeding animals high caloric diets, and that weight gain can be stopped by reducing caloric intake [185, 186]. Another study found that daily caloric intake was the main determinant of childhood obesity [187]. In contrast to calorie imbalance, the second hypothesis indicates that the type of carbohydrate eaten can cause obesity. There is evidence from studies on children that excessive fructose consumption may contribute to obesity regardless of overall energy intake [188]. Additionally, other studies show diets high in whole grains and fiber are negatively associated with adiposity [189]. These studies support the hypothesis that the type of carbohydrate eaten affects weight change and thus low-complexity carbohydrates could be causal to obesity. Our model gives a posterior mode score of 75.3, acknowledging that while evidence for both hypotheses exist, about five times more papers support the calorie imbalance hypothesis than the low-quality carbohydrate hypothesis.

#### Time needed for KM-GPT-DCH to correctly choose a hypothesis

How much time needs to pass after a new hypothesis is presented before KM-GPT-DCH can determine if the newer or older hypothesis is more likely? For the hypotheses presented here, this period varies from 0-6 years after the first proposal. All these determination dates from KM-GPT-DCH are before the new hypothesis was either generally accepted correctly or rejected correctly by the scientific community and public. So, while KM-GPT-DCH is likely useful for speeding up the process of correct acceptance or rejection of a new hypothesis or statement, it may not be typically capable of effectively evaluating a brand-new hypothesis. It is probably more typical that some literature about the newer hypothesis **after** its initial publication needs to exist for KM-GPT-DCH to make a significant evaluation.

### Supplementary Methods

#### The KM-GPT-DCH algorithm additional notes

Here, we use the simpler KM (KinderMiner) algorithm [190] for the co-occurrence step, versus Serial KinderMiner. One could, in theory, use SKiM (Serial KinderMiner) [2] to retrieve co-occurring documents of interest and then still perform the GPT (Generative Pretrained Transformer) step [4] and DCH (Direct Comparison of Hypotheses) analysis on these documents. This could be useful if one wanted to generate hypotheses and then test competing generated hypotheses.

#### Example KM-GPT-DCH prompt

The following is the prompt input to the LLM in the KM-GPT-DCH approach:

Biomedical Abstracts (verbatim):

PMID: XXX

Title: XX

Abstract: XX

===END OF ABSTRACT===

Available PMIDs for citation: XXX

Task:

Compare Hypothesis 1 vs Hypothesis 2 using ONLY the abstracts above that mention terms relevant to either hypothesis.

Classify each abstract, produce tallies, assign a continuous 0\100 score, and choose a decision.

Hypothesis 1:

Peptic ulcer is caused by bacterial infection.

Hypothesis 2:

Peptic ulcer is caused by psychological stress.

Continuous Scoring Guidelines (0\100):

**0:** Hypothesis 1 is **strongly refuted** by overwhelming and consistent evidence indicating that peptic ulcer is caused by bacterial infection. **Directly contradicts** the proposed outcome, while Hypothesis 2 is **strongly supported** by overwhelming and consistent evidence indicating that peptic ulcer is caused by psychological stress.

**Directly aligns with** the proposed outcome.

**25:** Hypothesis 2 (peptic ulcer is caused by psychological stress.) is three times more likely than Hypothesis 1 (peptic ulcer is caused by bacterial infection.).

**50:** Hypothesis 1 (peptic ulcer is caused by bacterial infection.) is equally likely compared to Hypothesis 2 (peptic ulcer is caused by psychological stress.).

**75:** Hypothesis 1 (peptic ulcer is caused by bacterial infection.) is three times more likely than Hypothesis 2 (peptic ulcer is caused by psychological stress.).

**100:** Hypothesis 1 is **strongly supported** by overwhelming and consistent evidence indicating that peptic ulcer is caused by bacterial infection. **Directly aligns with** the proposed outcome, while Hypothesis 2 is **strongly refuted** by overwhelming and consistent evidence indicating that peptic ulcer is caused by psychological stress. **Directly contradicts** the proposed outcome.

Guidelines for Intermediate Scores:

- **Proportional Scoring:** Assign scores between the defined anchor points based on the relative strength and consistency of the evidence. For example, a score of +66 would indicate approximately twice as much support for Hypothesis 1 compared to Hypothesis 2. In the case where there is no evidence for one of the Hypotheses, but there is supporting evidence for the other hypothesis, then this should generate a score in favor of the other hypothesis, the score value being dependent on the actual level of support for the other hypothesis. For instance, if there is no evidence at all for Hypothesis 2, and there are 10 abstracts that exhibit reasonably strong support for Hypothesis 1 (but do not provide absolutely unequivocal support for Hypothesis 1), then this should generate a score strongly in favor of Hypothesis 1, but not necessarily the max score of 100.
- **Avoid Clustering:** Do not disproportionately favor specific integer values unless the evidence explicitly warrants it. Ensure that scores are distributed across the range to reflect the nuanced degree of support or refutation.
- **Evidence-Based Justification:** Base the score on the cumulative evidence from the abstracts. Consider the number of abstracts supporting or refuting each hypothesis, the quality of the evidence, and the presence of any contradictory information.
- **Neutrality and Insufficiency:** A score close to **50** should be assigned when the evidence is inconclusive, mixed, or insufficient to determine the support of both hypotheses. Note that in the case where there is no evidence for Hypothesis 1 and no evidence for Hypothesis 2, this should generate a score of 50.
- **Consistent Reasoning:** Ensure that the reasoning provided for the score directly correlates with the assigned value, clearly demonstrating how the evidence leads to that specific score.

Best Practices to Maintain Objectivity:

- **Clear Definitions:** Utilize the anchor points to anchor your scoring decisions, ensuring each score has a clear and objective basis.
- Comprehensive Review: Thoroughly analyze all provided abstracts to capture the full scope of evidence before assigning a score.
- Unit Consistency: Keep the scoring consistent across different hypotheses by adhering strictly to these guidelines, avoiding subjective or arbitrary score assignments.

By following these continuous scoring guidelines, you can provide a nuanced and objective assessment of the support level for Hypothesis 1 compared to Hypothesis 2 based on the hypotheses that peptic ulcer is caused by bacterial infection and peptic ulcer is caused by psychological stress.

Output policy:

Return ONLY a single JSON object matching the schema below, inside a ```json code block.

JSON schema (for reference; do not print this schema):

{'type': 'object', 'additionalProperties': False, 'properties': {'per_abstract': {'type': 'array', 'items': {'type': 'object', 'additionalProperties': False, 'properties': {'pmid': {'type': 'string', 'pattern': '^[0-9]+$'}, 'label': {'type': 'string', 'enum': ['supports_H1', 'supports_H2', 'both', 'neither', 'inconclusive']}, 'evidence': {'type': 'array', 'items': {'type': 'string', 'maxLength': 300}}}, 'required': ['pmid', 'label']}}, 'score_rationale': {'type': 'array', 'items': {'type': 'string', 'maxLength': 1000, 'description': \Evidence-based rationale with PMIDs, e.g., 'Two RCTs report X (PMID: 123, 456)' that uses the scoring guidelines to justify the score.\}, 'minItems': 1, 'maxItems': 6}, 'tallies': {'type': 'object', 'additionalProperties': False, 'properties': {'support_H1': {'type': 'integer', 'minimum': 0}, 'support_H2': {'type': 'integer', 'minimum': 0}, 'both': {'type': 'integer', 'minimum': 0}, 'neither_or_inconclusive': {'type': 'integer', 'minimum': 0}}, 'required': ['support_H1', 'support_H2', 'both', 'neither_or_inconclusive']}, 'score': {'type': 'number', 'minimum': 0, 'maximum': 100}, 'decision': {'type': 'string', 'enum': ['H1', 'H2', 'tie', 'insufficient_evidence']}}, 'required': ['per_abstract', 'score_rationale', 'tallies', 'score', 'decision']}

#### Determining the acceptance/refutation date for the hypotheses

Acceptance and refutation dates for the hypotheses discussed in this paper come from review of the literature and other available data. Acceptance by the scientific community is based on the following literature. For the case of HPV (human papillomavirus) and cervical cancer, the 1996 National Institute of Health (NIH) conference where HPV was accepted as a leading cause of cervical cancer is the most relevant piece of information [191]. For the case of Scrapie being caused by prions (protein infection), the acceptance date of 1997 is supported by this paper [192]. For the case of bacterial infection being a leading cause of peptic ulcer, the acceptance date of 1994 is supported by the NIH conference in that year when it was concluded that antibiotic treatment is crucial for treatment of peptic ulcer [193]. For the case of vaccines causing autism, that was refuted by the Lancet retraction in 2010 [194]. For the case of hormone replacement therapy (HRT) being useful for prevention of cardiovascular disease, this was refuted by a large clinical trial in 2002 [41].

General public acceptance/refutation is difficult to assess but typically lags behind scientific acceptance. For the case of HPV causing cervical cancer, even as late as 2005, less than 20% of U.S. women were aware that HPV causes cervical cancer [195]. In general, men are less aware about HPV and cervical cancer [196]. General public awareness increased with the release of the Gardisil vaccine in 2006 [197], but awareness of the link between HPV and cervical cancer was still low as of 2007 with only 2.5% of British women aware that HPV causes cervical cancer [198]. Thus, we chose 2006 as a community acceptance date for HPV and cervical cancer, but the actual public awareness at this point was probably still low.

For the case of prions being infectious agents that cause diseases like scrapie, as of 2003, only 40% of university students were aware that prions cause mad cow disease [199]. General public awareness of the infectious agent being a non-living protein remains low as indicated by surveys of high-stakes individuals (hunters) in 2020-2022 indicate that only 21-38% of respondents were aware that prions cause chronic wasting disease (CWD) [200]. We set the public acceptance date as 2020+ to indicate that even as of 2020 or later many high-stakes individuals still were unaware that prions cause CWD. For the case of a bacterium causing peptic ulcer, as of 1997 only 27% of the U.S. public understood that a bacterial infection causes peptic ulcer [201]. It is difficult to find accurate surveys in the remaining several years. A 2012 study found low awareness of *H. pylori* (21-35%) amongst Chinese migrant workers [202]. However a 2024 study in which more than 50% had a college degree suggested that a high percentage (82.5%) of survey participants in Riyadh, Saudi Arabia were aware that *H. pylori* can cause peptic ulcer [203]. Thus, the acceptance rate will probably depend on education, socioeconomic status, and many other factors. We conservatively set the public acceptance date for the hypothesis that bacterial infection is a cause of peptic ulcer as 1997+.

For the case of the incorrect claim that vaccines cause autism, the number of people that think vaccines cause autism is increasing in the U.S. A 2020 U.S. Gallup poll showed that 10% believe vaccines cause autism, and 46% are unsure if vaccines cause autism or not [204]. These percentages increase in the 2024 U.S. Gallup poll which showed 13% of people believe vaccines cause autism, and 51% are unsure [205]. Thus, we do not set the public refutation date for the claim of vaccines causing autism as it is still unknown.

For the case of the incorrect claim the HRT would be useful for preventing ASCVD in peri- or post-menopausal women, after the publication of the Women’s Health Initiative study in 2002 [41], there was substantial media coverage, but this resulted in only moderate understanding by the public with 29% of women being aware of the WHI results by 2004 [206]. However, the response by clinicians was quick and substantial with the number of hormone therapy claims dropping by ~30% by the end of 2002. Claims continued to drop by 70% by 2009 [207]. Of course, clinicians may not be representative of the public. Based on the information above, we set the public refutation date for HRT as a preventive treatment for ASCVD to be 2004+.

See **Table 1** for scientific acceptance dates for historical hypotheses, and **Supplemental Table 1** for scientific and public acceptance dates for historical hypotheses.

#### Choosing A and B terms

Note that because KM uses an exact text matching search method, it is important to choose A and B terms to broadly capture the abstracts of interest. The user can choose multiple versions of terms using the pipe symbol (*e.g.* peptic ulcer| gastric ulcer). Care must be taken to not include synonyms that might be used in other contexts; however, the relevance filter will likely remove such abstracts before consideration by the frontier LLM. The number of abstracts retrieved and resulting analysis will depend on the A and B terms chosen. Because of the random sampling, if there is a large A-B abstract pool, different abstracts would be retrieved for different iterations (up to 50 per iteration), resulting in score variation. It is therefore recommended to run multiple iterations especially for a large pool of abstracts.

#### Estimating LLM Scoring Variability

If an identical set of abstracts is provided to an LLM repeatedly, the LLM can produce different scores. LLM scoring variance was quantified by re-running the same set of abstracts with the same set of hypotheses for 50 iterations and then calculating the variance of the LLM scores. We chose 20 different sets of abstracts and hypotheses to test, including two time periods from the peptic ulcer hypotheses (bacteria cause versus stress cause) starting in 1987 and in 2025, and two time periods from the cervical cancer hypotheses (HPV cause versus HSV cause) starting in 1983 and in 2025 (all with a 5 year window for getting abstracts). For peptic ulcer, 1987 was chosen because it is a year consisting of mixed findings (both bacteria and stress causes are supported) while 2025 is a year of mostly support for a bacteria cause. Similarly for cervical cancer, 1983 was chosen because it is a year of mixed findings supporting both HPV and HSV, while in 2025 mostly HPV is supported. Another 16 hypotheses with a range of scores were chosen from the 20 new controversial hypotheses, representing years 2022-2026 (see **Supplementary Data 3** for runs and scores). The variance of the scores on the $[0, 1]$ scale (${\sigma_{llm}}^{2}$ ) was computed for each run.

The Bernoulli variance, $\theta\left( 1-\theta\right)$, was fit to the variance of the scores (${\sigma_{llm}}^{2}$) with a linear model (see **Supplementary Figure 3**). Because $\theta$ is unknown, the average LLM score on the [0, 1] scale across the 50 iterations was used in place of $\theta$. For the linear model, the y-intercept was set to zero because the variance cannot be less than zero.

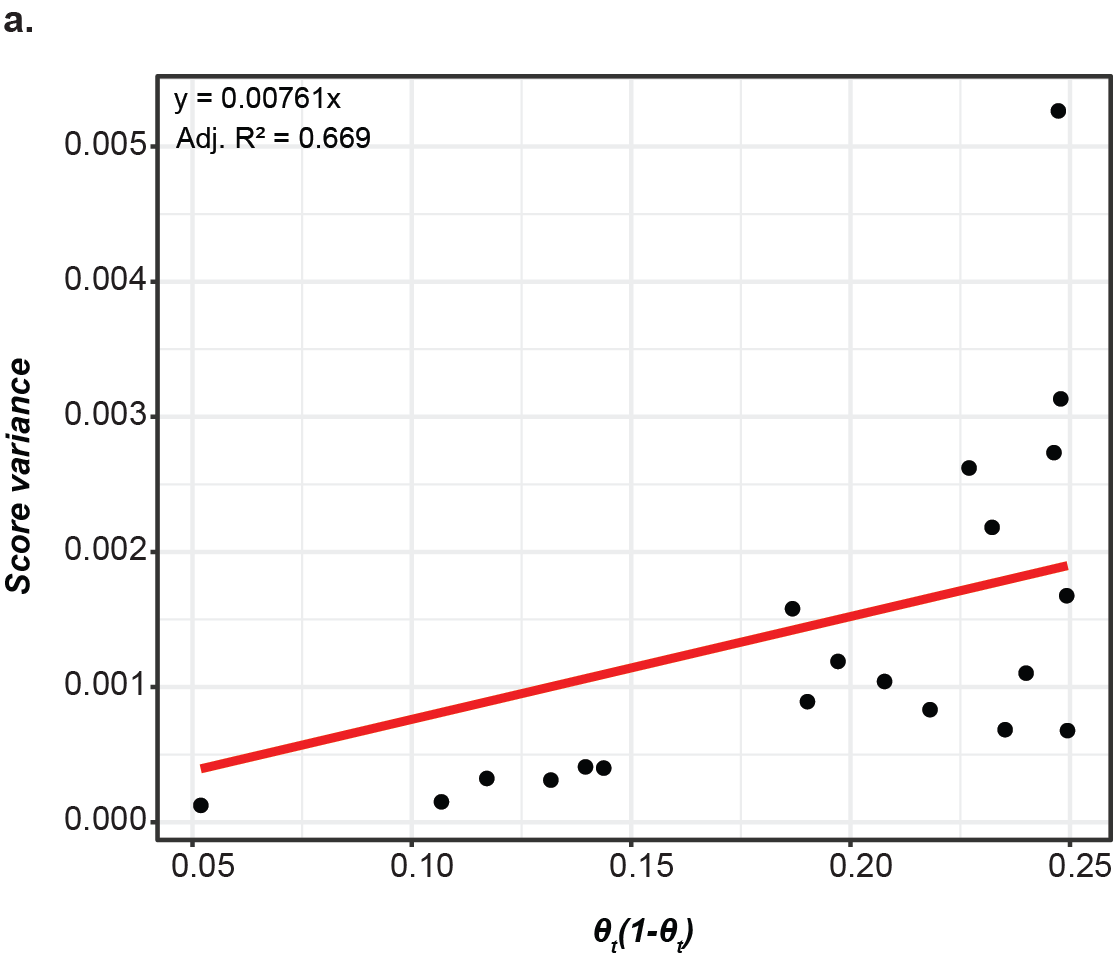

**Supplementary Figure 3. LLM score variance vs. Bernoulli variance (**$\boldsymbol{\theta}\left( \mathbf{1-}\boldsymbol{\theta} \right)$**) from abstract sampling**

The figure shows the slope of the line between the LLM score variance (${\sigma_{llm}}^{2}$) on the y-axis and the Bernoullis variance ($\theta\left( 1-\theta\right)$) on the y-axis, where the slope is found by $y = mx$and the y-intercept is set to zero.

### Other Supplementary files

**Supplementary Table 1:** Acceptance/ Refutation dates of Historical Hypotheses and KM-GPT-DCH resolution dates.

**Supplementary Table 2:** Summary of reference evaluations by LLM-interfaces for pre-study time cut-offs (sheet1) and post-study time cut-offs (sheet 2).

**Supplementary Table 3:** Summary of reference evaluations by Robin, a multi-agent system for literature review using Crow interface.

**Supplementary Table 4:** Summary of current controversial or competing hypotheses that were evaluated using KM-GPT-DCH.

**Supplementary Data 1**: ***Detailed Q and A web-based LLMs.*** All questions and answers posed to and given by the chat-based LLMs and Robin for evaluation that are summarized in Supplementary Tables 2 and 3.

**Supplementary Data 2: *All KM-GPT-DCH outputs.*** All responses for every time point and iteration from KM-GPT-DCH for each set of hypotheses discussed. Zenodo DOI: **10.5281/zenodo.20547913.**

**Supplementary Data 3**: All results from the comparative hypotheses run for variance testing.
